# Genetic Structuring and Conservation of Asian Sockeye Salmon: Identification of Regional Stock Complexes

**DOI:** 10.1101/2024.06.01.596976

**Authors:** Anastasia M. Khrustaleva

## Abstract

In order to describe large-scale spatial structure of Asian sockeye salmon the variability of 45 SNP loci was analyzed in 22 samples from the North-West coast of the Pacific Ocean. Three large regional population complexes were identified: southwestern Kamchatka, Kamchatka River basin, and the North-East (comprising stocks from Koryak Highlands). Populations within the identified complexes are connected by gene migration and have a common origin, close geographic proximity, comparable climatic, landscape and environmental conditions in the freshwater and early marine periods of life. Populations confined to watersheds of the North coast of the Sea of Okhotsk (Palana and Okhota rivers), along with island populations, displayed noteworthy distinctions from the isolated population complexes. We hypothesize that the marked divergence observed in island populations is primarily caused by genetic drift occurring during long periods of isolation. The pronounced divergence of Palana River population may be the result of both genetic drift and natural selection, driven by the challenging smoltification and juvenile transition to the ocean, along with local adaptations during spawning and early life periods in the Palansky Lake. At the same time in the Okhota River population, demographic factors such as genetic drift and bottlenecks played a key role.

## Introduction

Sockeye salmon *Oncorhynchus nerka* (Walbaum) is a commercially important species that has been an object of both fishery and artificial breeding across its extensive distribution range, which encompasses the entire northern part of the Pacific Rim (Burgner, 1991). It is most numerous in the North America, where 80-85% of its stock are reproduced, while Asian populations contribute to about 15-20% (Forrester & C.R., 1987; Bugaev, 2011). Sockeye salmon has the most complex population structure among all species within the genus *Oncorhynchus*. Large populations of Pacific salmon display metapopulation characteristics (Schtickzelle & Quinn, 2007). This implies that they comprise a system of relatively isolated populations interconnected through minimal individual migrations and capable to extinction and subsequent recolonization at the expense of other components of the system. Such dynamics contribute to the overall stability of the system over successive generations. In general, populations of sockeye salmon from different rivers are subdivided into distinct local subpopulations and seasonal races, which spawn during the summer (early form) and autumn (late form) (Burgner, 1991; Quinn, 2005). Apart from temporal and geographic intraspecific units, sockeye salmon exhibit a life-history dichotomy in their freshwater rearing environments (life-history ecotypes) (Quinn, 2005; Wood et al., 2008): lake-type populations rear in lakes for one to three years (more often two years) before travelling to the ocean to feed whereas sea/river populations rear in river habitats normally for one year, or even less (Pavey et al., 2010). Foraging, water current, and predation differ significantly between the lake-type and riverine populations affecting stable morphological and genetic differences arising as a consequence of local adaptations and isolation by adaptation (Lin et al., 2008). For instance the river ecotype has a deeper, shorter caudal peduncle associated with swimming against a current and a deeper body, whereas lake-type sockeye salmon have a more streamlined body shape (Pavey et al., 2010). Furthermore, sockeye salmon has high-level organization of population systems, such as large regional population complexes (Varnavskaya, 2006) or eco-geographic units (EGUs) (Zhivotovsky, 2016). Although both concepts are largely similar, the first one has a slightly broader scope, and we will adopt it for future use. Such complexes have their own biological, ecological, demographic and genetic characteristics and diverse local adaptations to a variety of environmental conditions, which allows the entire system to survive with significant environmental changes and anthropogenic pressure due to the restoration of endangered components at the expense of other populations of the same complex (Hilborn et al., 2003). These complexes are formed due to the shared descent, robust connectivity through migratory patterns, congruent adaptations arising from similar ecological and physical-geographical habitat attributes, and stabilization driven by ecological diversification (Olsen et al., 2008). The sustainability of population systems (both metapopulations and regional complexes) of sockeye salmon is maintained by their inherent biocomplexity. Within the context of Pacific salmon, biocomplexity is an interconnected network of local populations that as a whole maintain a relatively constant overall stock productivity, achieved through diverse components and a range of life strategies (Hilborn et al., 2003).

Since 2017, in a number of watersheds of Kamchatka, significant changes in the dynamics of the spawning run, biological indicators, and intrapopulation diversity of sockeye salmon have been observed. These changes have arisen as a consequence of density-dependent, trophic, and both local and global climatic and hydrological factors. However, the primary driver has been an imbalanced pressure on the population system components stemming from unsustainable management practices and non-selective fishery (Lepskaya et al., 2017; “Information …,” 2019; Koval et al., 2020). The main reason is the insufficiency or ignorance of information regarding the population structure, ecological dynamics, and temporal differentiation of salmon populations during the in the planning and organization of fishing. The organization of sustainable fishery, coupled with the protection, artificial propagation, and management of salmon populations, requires an extensive scientific base. This base should encompass fundamental aspects such as the delineation of exploited stock boundaries, assessment of biocomplexity, genetic diversity, adaptive capabilities, and constraints. It should also encompass considerations of conservation prospects, especially in scenarios involving intensive artificial reproduction and opportunities for recolonization in the event of overexploitation. It is also important to assess the background for successful recolonization, evaluate the potential of donor populations, and gauge the reproductive success of migrating individuals. A comprehensive, large-scale investigation into the population structure of Asian sockeye salmon, involving the identification of regional population complexes, their differentiation, and an exploration of their origin and historical development, will provide valuable insights into the demographic, ecological, and evolutionary processes occurring within these systems. Such a study has the potential to address, if not fully answer, many of the questions that have been raised in this context.

Over 95% of Asian sockeye salmon stocks are concentrated within the Kamchatka Peninsula, with primary reproduction centers in the Kamchatka (East coast) and Kurilskoe Lake, Ozernaya (South-West coast) river basins. In these watersheds, about 80–90% of the total catch of the species in the Russian Far East is annually caught (Bugaev, 2011). Among secondary sockeye salmon stocks, several distinct populations have a relatively high abundance: on the west coast it’s the populations of the Bolshaya and Palana rivers, in the east of Kamchatka – Apuka R. and Pakhacha R. populations (Shubkin & Bugaev, 2023). Chukotka is the second most important region of sockeye salmon reproduction in Asia after the Kamchatka peninsula. In the watersheds of Eastern Chukotka, the largest population of sockeye salmon inhabit the Meinypylgin lake-river system (Golub, 2003). On the North Okhotsk coast, sockeye salmon is not numerous. The largest stock of the Okhotsk region is reproduced in the basin of the Okhota river (Nikulin, 1975). However, it is not large and has no significant commercial value (Chereshnev, 2008). Relatively small populations of sockeye salmon inhabit watersheds of the Commander Islands, Hokkaido Island and the Kuril Islands − Shumshu, Paramushir, Urup and Iturup (Shedko, 2002; Zhulkov et al., 2012). Despite their limited commercial significance and relatively small numbers, island populations of sockeye salmon are of exceptional scientific interest and may provide important insights into the origins and evolutionary history of sockeye salmon in Asia. Moreover, these island populations are a convenient model for studying microevolutionary processes and potential pathways of adaptive evolution within the species. Island populations frequently exhibit unique ecological, morphological, and genetic characteristics, a phenomenon known as the ’island syndrome’, which is formed as a result of historical and demographic processes and under the influence of climatic, orographic, ethological and biocenological factors in conditions of isolation (Baeckens & Van Damme, 2020).

The distinctive characteristics that make the island’s biodiversity so special also make it particularly susceptible and vulnerable. The level of diversity in island populations is usually low, and besides, their numbers are not large, all this makes them more prone to extinction. Furthermore, due to restricted resources, reduced abilities to disperse, and residence in relatively stable and predictable marine climates, island populations, often located at range edges, are more vulnerable to limiting factors. They evolve survival strategies developing local adaptations, interdependence, co-evolution, and mutual influence within limited list of species and factors, rather than broad defense mechanisms against a wide array of predators, competitors, infections, and variable physical environmental conditions. Consequently, a disproportionately high number of species extinctions have been recorded on islands compared to continental systems. Island populations are exceptionally vulnerable to anthropogenic impacts and demand focused efforts and specific conservation measures. Therefore, it is of the utmost importance to identify and assess complexes of island populations to determine their conservation potential.

Asian sockeye salmon island populations remain inadequately explored due to their limited numbers, logistical challenges due to the remote and inaccessible locations, and the complexities associated with organizing fisheries in uninhabited areas. However, between 2006 and 2014, we were able to collect samples from several sockeye salmon nursery lakes on the Kuril and Commander Islands. The study of the polymorphism of the mtDNA control region showed that the sockeye salmon from the lakes of Iturup Island was characterized by moderate haplotype and nucleotide diversity, while the genetic diversity of the sockeye salmon from the Northern Kuriles and Bering Island was significantly lower compared to the continental populations (Khrustaleva, 2016; Khrustaleva et al., 2020). Furthermore, unique haplotypes were found within the South Kuril sockeye salmon populations, which are transitional forms between two Asian mtDNA haplogroups. We believe that these populations of sockeye salmon may have originated from the southern refugium that existed during the last Pleistocene regressions in the Hokkaido region and may have avoided secondary contact during the last wave of colonization of the Asian coast in the early Holocene.

Given the limited number of marker-based genetic studies on Asian sockeye salmon, our primary objective is to conduct a comprehensive investigation into the population structure of this species on a regional scale, encompassing its entire distribution range along the West Coast of the Pacific Ocean. We will place particular emphasis on studying island populations, which face heightened vulnerability due to global climate change in the North Pacific and escalating anthropogenic pressures. Additionally, we aim to identify regional population complexes that can act as protective buffers for these island populations, helping to mitigate the negative impact of environmental changes or unsustainable commercial human activities. Finally, our research will explore the ecological, genetic, and historical factors that contribute to the distinctions observed among the complexes and populations of Asian sockeye salmon.

## Materials and Methods

### Study area and sample collection

The samples were collected in 2003 through 2008 in the rivers of the East and West coasts of Kamchatka peninsula, Chukotka peninsula, mainland coast of the Sea of Okhotsk, Kuril and Commander Islands (Table 1, Figure 1). Sockeye salmon adults were caught using river seine nets in the river beds and lake creeks at a distance of 5-30 km from the river mouth during the mass run of sockeye salmon, as well as directly in the spawning lakes (Supplementary Table S1). Most Kamchatka samples were obtained from fishing companies directly after catch in local fisheries. In the Bolshaya River, smelt juvenile fishes were caught using minnow seine in the upper reach of Plotnikov River and in the lower course of Bystraya River (Supplementary Table S1, Figure 1). The pectoral fin and liver tissue samples were fixed in 96% ethanol.

**Fig. 1.**
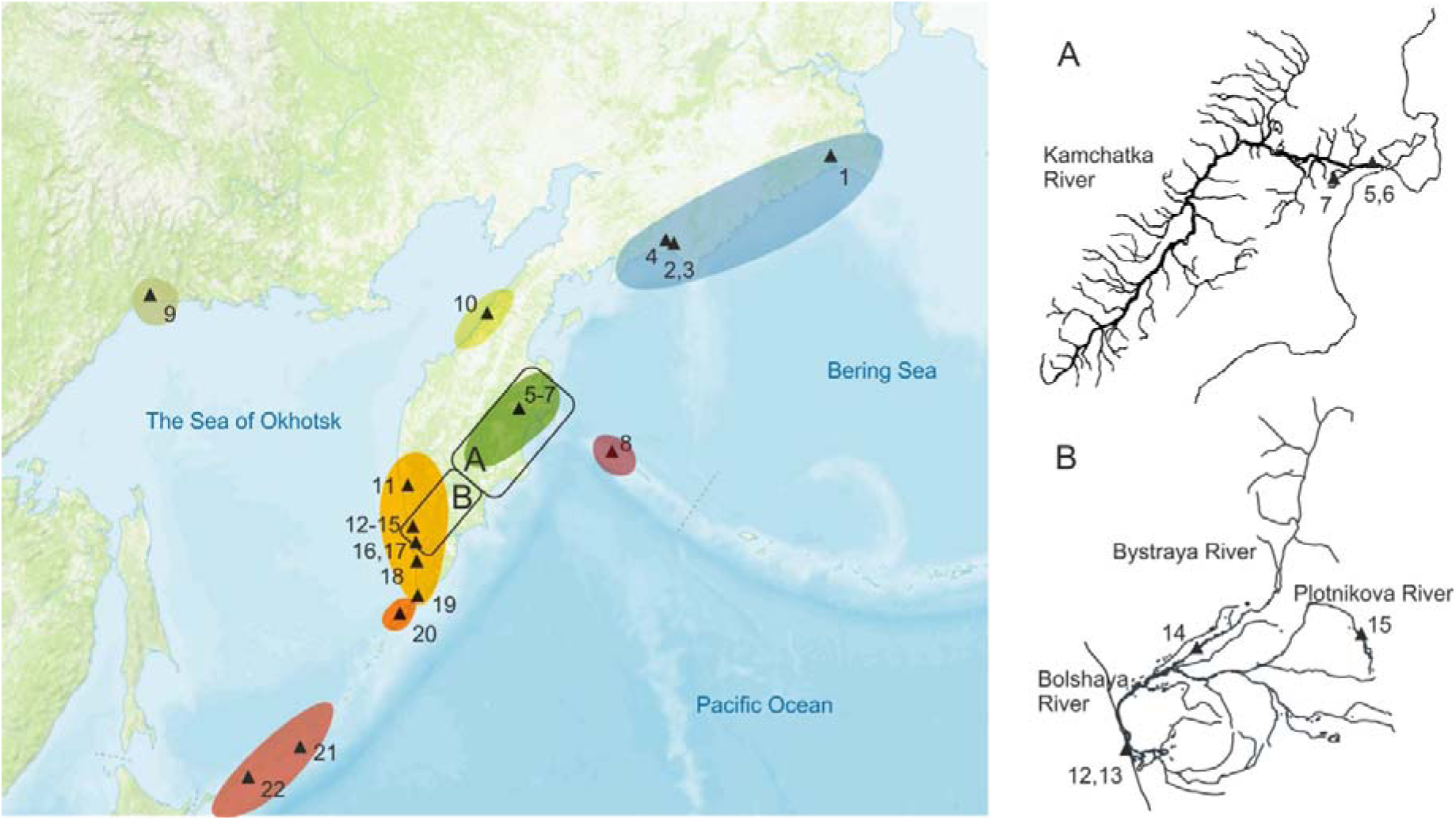
Schematic map of the study area with sampling points (triangles). The point’s annotations are given in Table. 1. Regional complexes of Asian sockeye salmon are marked with different colors.

**Table 1.**
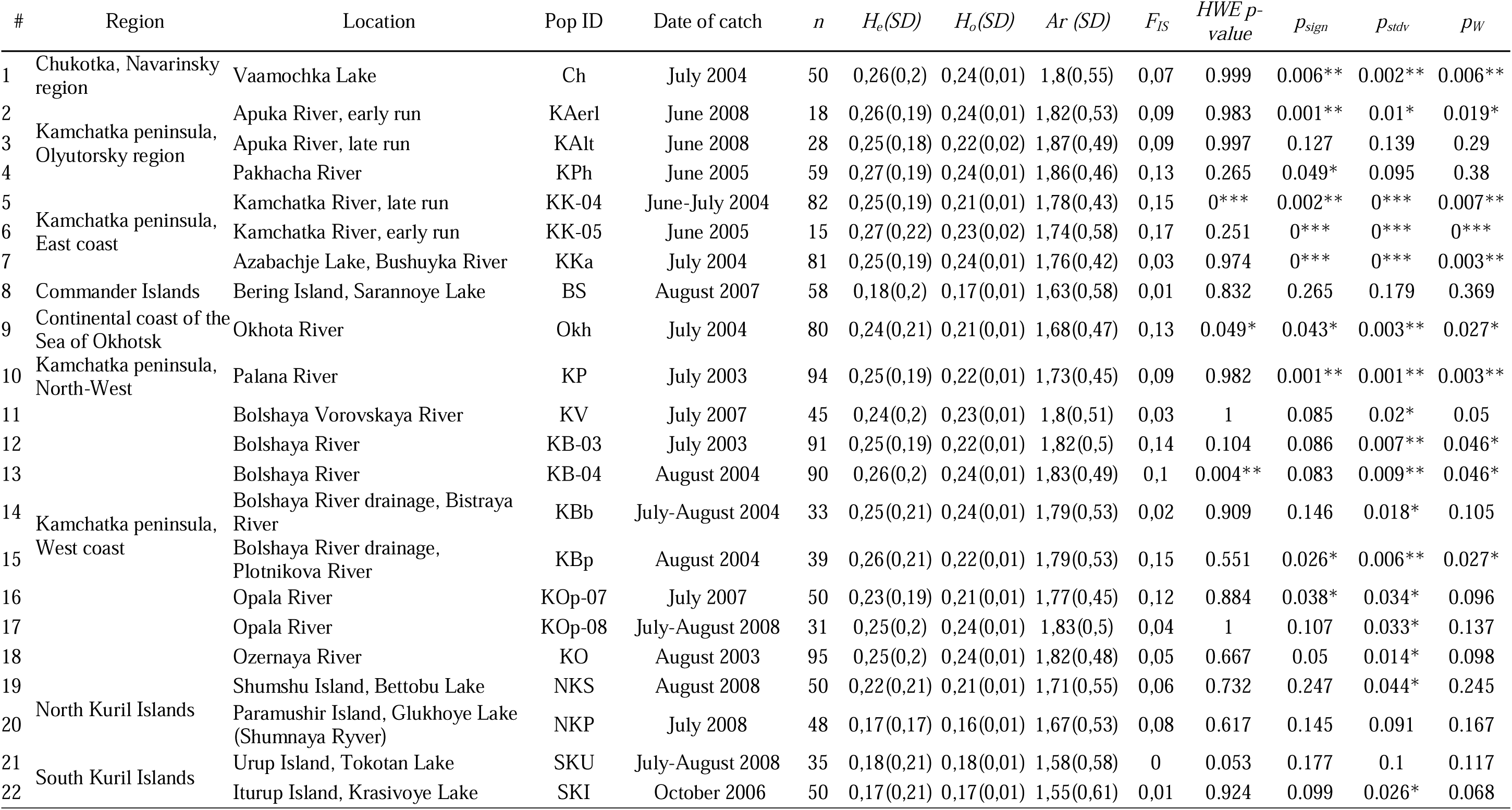
Samples characteristics, regions and locations, population IDs, date of catch, and summary statistics for 40 SNP loci: mean expected (*He*) and observed (*Ho*) heterozygosities, allelic richness (*Ar*), the inbreeding coefficients (*F_IS_*), the results of the exact tests on Hardy–Weinberg equilibrium (*HWE p-value*), and the results of sign test (*p_sign_*), standardized differences test (*p_stdv_*) and two-tail Wilcoxon sign-rank tests (*p_W_*) for heterozygote excess, * − *p* < 0.05, ** − *p* < 0.01, *** − *p* < 0.001.

### SNP genotyping and data borrowing

Genomic DNA was extracted with Qiagen DNeasy 96 tissue kits (Qiagen, Valencia, California). TaqMan-PCR using Fluidigm 96.96 Dynamic Arrays (Fluidigm, San Francisco, California) allowed for the genotyping of 95 individuals per 96-well plate (with one inlet used as a no-template control using tris-EDTA buffer) was carried out following the protocol of Seeb et al. (Seeb et al., 2009). All individuals (n = 1226) were genotyped for 45 SNP loci (Habicht et al., 2010) including three mitochondrial and 42 nuclear loci localized in structural and regulatory genes, dispersed repeats, and EST (Supplementary Table S2). In total 22 sockeye salmon samples from 14 localities of the Asian coast of the Pacific Ocean were analyzed. For a more extensive analysis of the regional subdivision of Kamchatka sockeye salmon, open data on the same set of loci (http://www.tandfonline.com/doi/suppl/10.1577/T09-149.1?scroll=top) of Dr. C. Habicht and coauthors (Habicht et al. 2010) were used (Supplementary Table S3, Figure S1). Hereinafter, the *One_* prefix in loci names is omitted for brevity.

### Statistical analysis

Allelic frequencies, observed and expected heterozygosities (*Ho, He*) for each locus (excluding mtDNA SNPs), and estimates of the allelic richness (*Ar*) by rarefaction using the smallest sample size were obtained in hierfstat (Goudet, 2005). Deviations from Hardy-Weinberg expectation (HWE) were evaluated across all loci for each population by exact test (using the Markov chain (MC)) implemented in GENEPOP v4.0 (Raymond & Rousset, 1995). Default parameters were used for the MC algorithm (dememorization = 1,000; batches = 20; iterations per batch = 5,000). The Weir and Cockerham (1984) *F_ST_* and *F_IT_* statistics for each locus and global *F_ST_* statistics as well as exact G-tests for genic and genotypic differentiation were calculated using GENEPOP v4.0. Tests for linkage disequilibrium between all loci pairs were performed using simulated exact tests in GENEPOP v4.0. Benjamini-Hochberg FDR correction was applied for all multiple tests. The necessary condition for pooling of putatively linked loci was established as in (Habicht et al., 2010): if tests for linkage disequilibrium are significant in more than half of the samples, then either the less informative locus from the pair is dropped or the two loci is combined into a composite genotype. All three mtDNA SNPs were combined into a composite haplotypes for baseline evaluation (*Cytb_CO*). In order to select a set of neutral loci, a panel of 45 SNP markers was iteratively screened to identify outlier loci at different spatial scales, either analysing all population samples together or evaluating various combinations of samples from different locations (Supplementary Figure S2). Outlier detection was performed using coalescent simulations under the hierarchical island model to obtain p-values of the locus-specific F-statistic conditioned on observed levels of heterozygosities implemented in in Arlequin 3.5 (Excoffier & Lischer, 2010). The isolation by distance hypothesis was tested using the Mantel-tests for putatively neutral loci in the ade4 R package (Dray & Dufour, 2007). The evidence of population “bottlenecks” was identified using the Bottleneck 1.2.02 (Cristescu et al., 2010). This analysis is based on the loss of rare alleles predicted in recently bottlenecked populations, resulting in heterozygosity excess. As this method assumes that markers are selectively neutral, we only used non-outlier loci. We used the infinite alleles model (IAM) as the most appropriate evolutionary model for SNP loci. To test for significant heterozygosity excess compared to the level predicted under mutation-drift equilibrium, we used three tests: a sign test, a standardized differences test, and a one-tailed Wilcoxon signed rank test. A bottleneck was considered verified if all three tests were significant.

The Cavalli-Sforza chord distances evaluation and reconstruction of trees by the Neighbors Joining (NJ) method were carried out in Rphylip package (Felsenstein, 1989). Based on the distance matrices and the obtained bootstrap estimates (1000 iterations), a phylogenetic networks were built using the Neighbor-Net (NN) algorithm in the phangorn R package (Schliep, 2011). Principal coordinate analysis (PCoA) based on Euclidian distances between individual genotypes was carried out in hierfstat (Goudet, 2005). Principal component analysis (PCA) was performed using the R libraries factoextra (Kassambara & Mundt, 2020) and FactoMineR (Lê et al., 2008). The discriminant analysis of principal components (DAPC) was performed using adegenet 1.3-1 R package (Jombart & Ahmed, 2011). The population structure was assessed de novo, pre-determining the number of clusters in the entire dataset using iterative analysis of k-means. The optimal number of clusters (K), or genetic groups, is often defined as the K with the lowest Bayesian criterion (BIC) values among the identified clusters. However, we opted to use the scree test (choosing a point on the curve of BIC dependence on the number K after which the average rate of change of the function is significantly reduced). The DAPC procedure then used the optimal number of groups and the first 32 identified principal components. On the next step, the probabilities of each genotype belonging to a given cluster were graphically visualized.

Further clustering analyses were completed with STRUCTURE (Pritchard et al., 2000) where three independent runs for each number of cluster − K (2–22) were conducted using the admixture model at 50,000 iterations with a burn-in of 10,000. The most probable number of population clusters was determined by the estimation of ΔK in STRUCTURE HARVESTER (Earl & vonHoldt, 2012). In the last step, we used GENELAND v.0.3 (Guillot et al., 2008), which incorporates geographic information (coordinates) in order to estimate the number of panmictic groups and locating their spatial boundaries. The model used was the correlated frequency model, and a posterior probability map at the optimal cluster number, K = 8, was built, using 100000 Markov chain Monte Carlo (MCMC) iterations, with a thinning interval of 10000. Data visualization was performed in R using ggplot2 library.

## Results

### Characteristics of loci and Diversity within Asian Sockeye Salmon Populations

Of the 45 loci analyzed, 43 were polymorphic in at least one sample, with the exception of *p53-576* and *RAG1-103*. The tests for linkage disequilibrium (LD) for the *MHC2_190v2* and *MHC2_251v2* loci were significant in seven out of 22 samples, which did not satisfy the linkage criterion by Habicht at al. (Habicht et al., 2010). Concurrently, these two substitutions are located within the *Onne-DAB* gene of the Major Histocompatibility Complex (MHC) at a distance of 62 base pairs from each other. This physical proximity strongly indicates linkage, prompting their analysis both independently and as a single linkage block or a locus, denoted as *MHC2*, with distinct allelic/genotypic variants referred to as haplotypes/genotypes. LD-tests also revealed a correlation between the genotypes of two pairs of loci *GPDH − GPDH2* and *GPH-414 − MHC2_251* in the sample from the Kamchatka River (2004) and the *zP3b − MARCKS-2* pair in the sample of the Bolshaya River (2003) (Fig. 2A). Nonetheless, it is evident that there is insufficient basis to their blocking. Thus, after exclusion of monomorphic and blocking of linked loci, the number of analyzed SNPs decreased to 40.

**Fig. 2.**
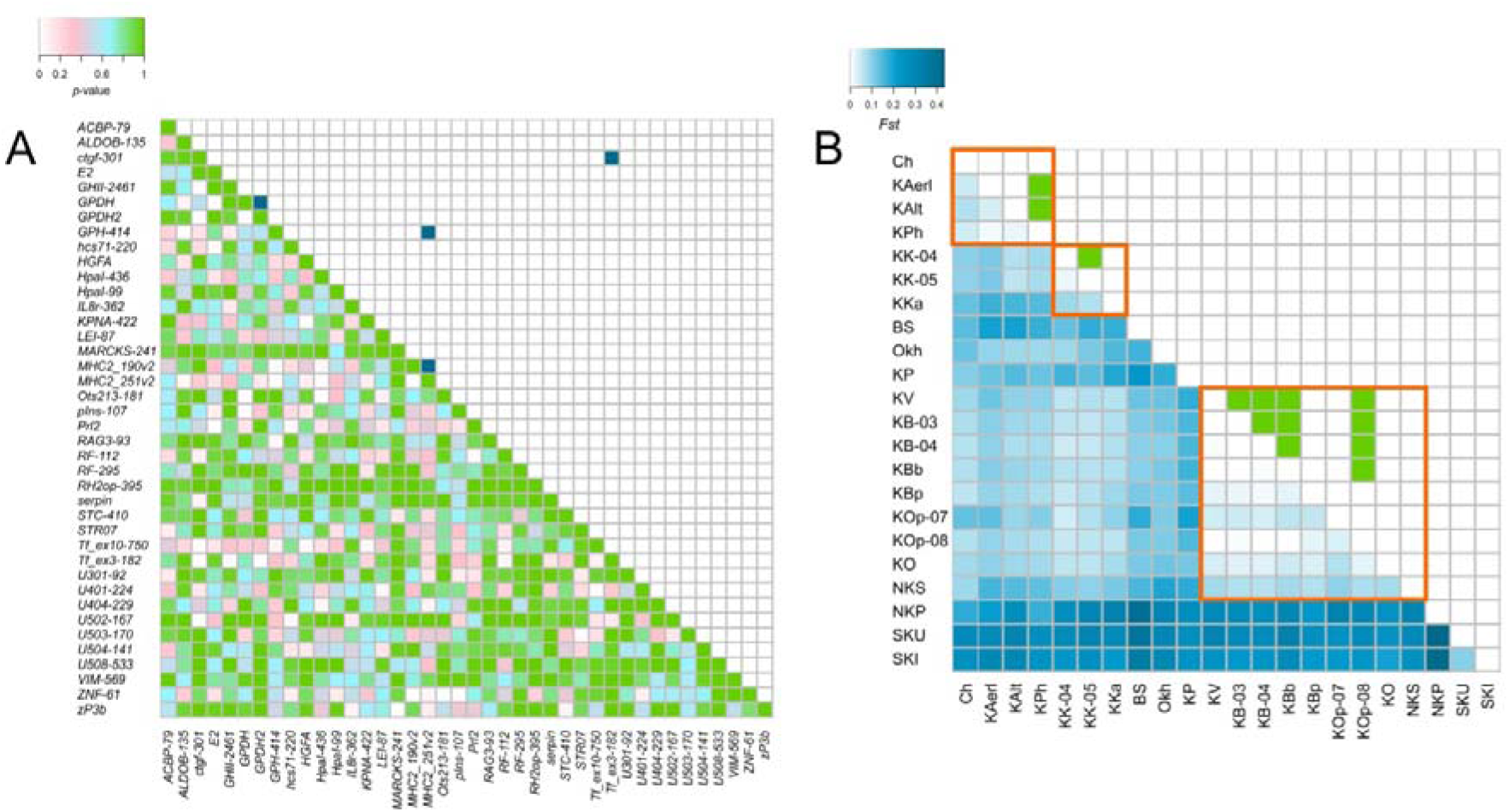
(**A**) The results of the LD-tests are presented as the heat map, below the diagonal − the significance levels (*p*-values) when testing the hypothesis about the genotypic equilibrium, above the diagonal − significant deviations from the equilibrium after FDR correction. Mitochondrial loci were excluded from this analysis; (**B**) Pairwise *F_ST_*matrix for Asian Pacific Coast sockeye salmon samples (below the diagonal). Above the diagonal − shaded cells indicate non-significant differences in paired exact-tests on genic differentiation after FDR correction. Sample annotations are given in Table. 1.

Estimates of intrapopulation genetic diversity in 22 samples of sockeye salmon are given in Table. 1, Supplementary Table. S3, and Figure S2. After FDR correction, a significant deviations from Hardy–Weinberg equilibrium was noted in samples from the mouth of the Bolshaya River (KB-03 and KB-04) at the *ZNF-61* locus (KB-03: *p* = 0.0004, *F_IS_* = 0.39; KB-04: *p* = 0.0002, *F_IS_* = 0.40) and in the sample from the Okhota River (Okh) for the *GPH-414* locus (*p* = 0.0001, *F_IS_*= 0.42). In addition, significant deviations from HWE expectations were revealed for the loci of the MHC2 complex: when they were considered separately, a significant deficit of heterozygotes for both loci was noted in the sample from the mouth of the Kamchatka River (KK-04) (*MHC2_190v2*: *p* = 0.0004, *F_IS_* = 0.65; *MHC2_251v2*: *p* = 0, *F_IS_* = 0.95) and for the *MHC2_251v2* locus in the sample from the Plotnikova River (KBp) (*p* = 0.0003, *F_IS_* = 0.61). In total, for all loci, tests for HWE were insignificant only in three samples after FDR-correction (Table 1).

### Population Differentiation: Understanding Genetic Variation and Structure

The heterogeneity of allelic and genotypic frequencies in all dataset was revealed by the results of G-test on genic and genotypic differentiation of populations. No significant inter-sample differences were found only for some loci − *ctgf-301, U502-167, MARCKS-241, Tf_ex3-182*, and *RH2op-395*. By the results of paired tests the alleles and genotypes frequencies of SNP loci did not differ significantly (after FDR correction) in samples from the rivers of the Koryak Highlands (KA and KPh), from the lower reaches of the Kamchatka River, and from watersheds of the South-East coast of the Kamchatka Peninsula (Fig. 2B).

The intersample genetic diversity, estimated by the *F_ST_* value, averaged 0.135 (*p* = 0). For individual loci it varied from 0.0009 (*MARCKS-241*) to 0.405 (*serpin*). According to the matrix of paired *F_ST_*, a pronounced differentiation of sockeye salmon from the South Kuril Islands and Paramushir Island, as well as the proximity of the populations of Southwestern Kamchatka, including the Shumshu Island, and the relatively high similarity of the populations of the Koryak Highlands (Ch, KA and KPh) are clearly traced (Fig. 2B).

### Clustering with PCoA, PCA, DAPC, and phylogenetic relationships of populations

As a reconnaissance study of the population structure of sockeye salmon in Asia, a PCoA analysis of individual genotypes of sockeye salmon from various localities of the Asian part of the range was carried out (Supplementary Figure S3). On the corresponding graph, it is difficult to delineate most of populations clusters, but individuals caught in the lake-river systems of the Southern and Northern Kuril Islands (samples SKI, SKU and NKP) and the North coast of the Sea of Okhotsk (samples KP and Okh) are well separated from the main point pool. To distinguish regional groups of populations, the samples were ordinated on the space of main principal components using the PCA (Fig. 3A). In sum, four main components were identified, explaining in total more than half of the variability (53.4%) of the genetic traits (allelic frequencies of 40 polymorphic SNP loci). The first component explains primarily the variability of the two loci – *serpin* and *HGFA*, as well as two mass haplotypes of the mtDNA locus; the second − *STC-410, ACBP-79, GPH-414* (Supplementary Figure S4).

**Fig. 3.**
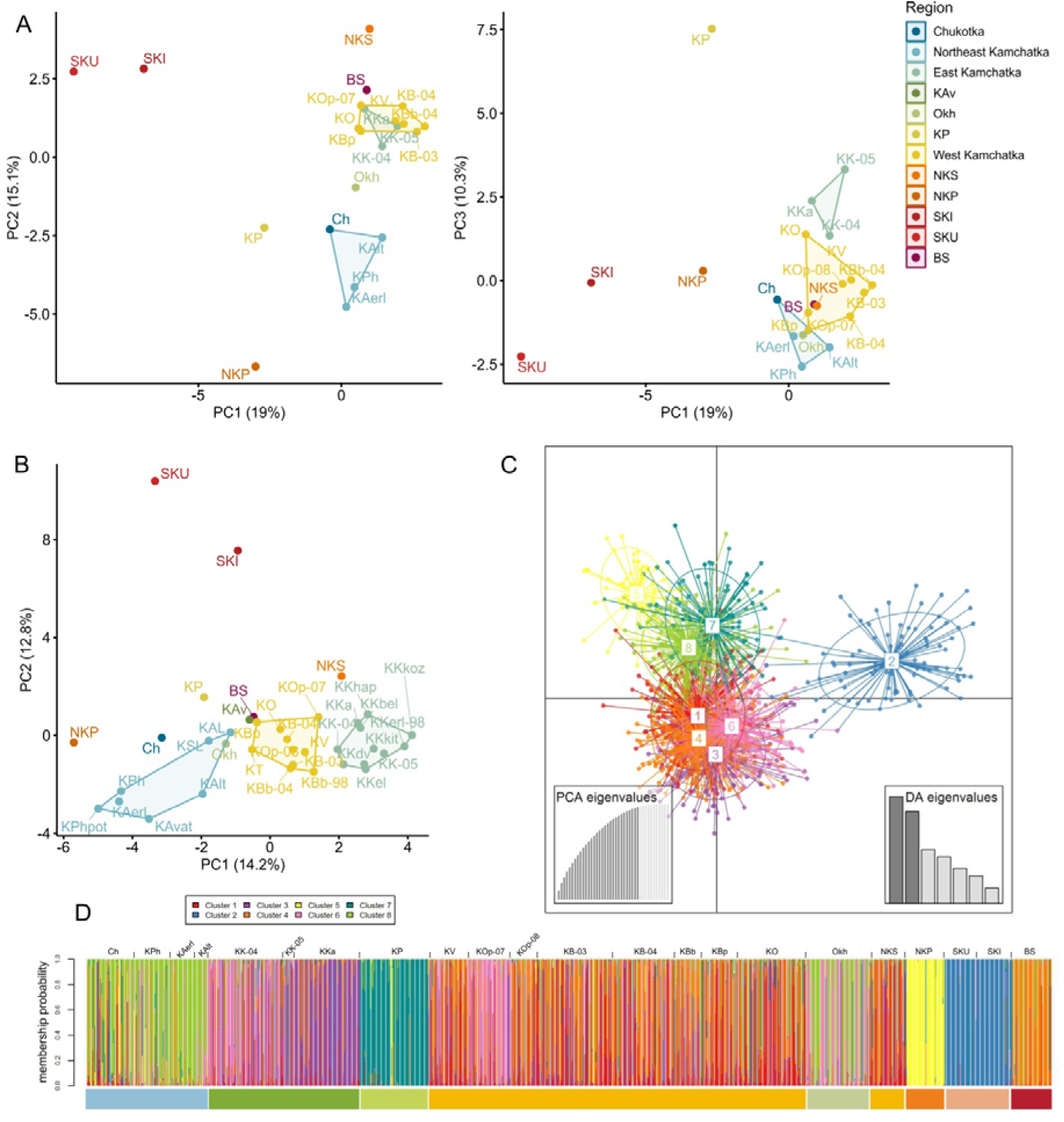
(**A**) PCA results showing population clustering based on the first two, and the first and the third principle components (PCs), own data; (**B**) The same, using our data and data from (Habicht et al., 2010). Sample annotations are given in Supplementary Table S3. (**C**) Clustering of the samples based on DAPC, clusters identified by the k-means method, the axes represent the first two linear discriminants; (**D**) An admixture analysis with k = 7 using DAPC, the histogram showing the probabilities of assigning individuals to the clusters. To indicate the regional affiliation of the samples, color bar is used, according to the color scheme in Fig. 1.

In the space of the first two principal components, the samples of sockeye salmon from western and eastern Kamchatka formed a dense cluster. Near the cluster there were located samples from the watersheds of the mainland coast of the Sea of Okhotsk (Okh) and Bering Island (BS). (Fig. 3A). Populations of the coasts of the North-East Kamchatka and Chukotka were isolated from this cluster along the second component. In addition, the Palana River sample was distant from the other populations of the West Kamchatka peninsula along the both components, whereas the third component differentiated the Palana sample to the greatest extent. The island populations of the North and South Kuril Islands were the most divergent from the others. (Fig. 3A). However, attempts to separate clusters corresponding to regional groupings of sockeye salmon were not successful. Furthermore, the proportion of genetic trait variability explained by the first pairs of components was relatively small, not exceeding 35%. This suggests that a significant percentage of genetic variability may be attributed to other factors requiring more scrupulous analysis. To achieve a higher level of detail, we conducted a PCA analysis that incorporated our own dataset, supplemented by data from (Habicht et al., 2010). By including thirteen samples from Kamchatka populations (Supplementary Table S3, Figure S1) we were able to differentiate large regional complexes of South-West Kamchatka (the South-West SW), Koryak Highlands including North-Eastern Kamchatka and Chukotka (the North-East, NE), as well as the Kamchatka river basin (KR) (Fig. 3B).

To classify individual genotypes and delineate the genetic subdivisions of sockeye salmon along the Asian Pacific coast, we conducted DAPC analysis employing a de novo clustering approach, enabled the identification of distinct groups (K=8) within the entire dataset (Supplementary Figure S5, Fig. 3C). Certain groups were represented by of individual populations: from Paramushir Island, Bering Island, as well as Palana and Okhota rivers, as depicted in Figure 3D. The mainland populations were divided into three large regional clusters corresponding to those identified earlier: SW (including the Shumshu Island), NE, and KR. Two samples from the South Kuril Islands were combined into a separate cluster (Fig. 3D).

To elucidate population relationships and the proximal chronological sequence of their divergence, we attempted to construct a split phylogenetic network. The original tree and the resulting network had a star-shaped topology and were divided into 2 clusters formed by Kamchatka peninsula populations on one side, and NE complex, the North Okhotsk sea coast (KP & Okh), and all island populations on the other (Fig. 4A). The network’s topology revealed four distinct clusters, corresponding to the population groupings the same as in PСA and DAPC tests. Notably, samples from the North Okhotsk sea coast and all island populations exhibited substantial divergence. The low bootstrap support observed in the source tree underscores the inherent instability of the net topology.

**Fig. 4.**
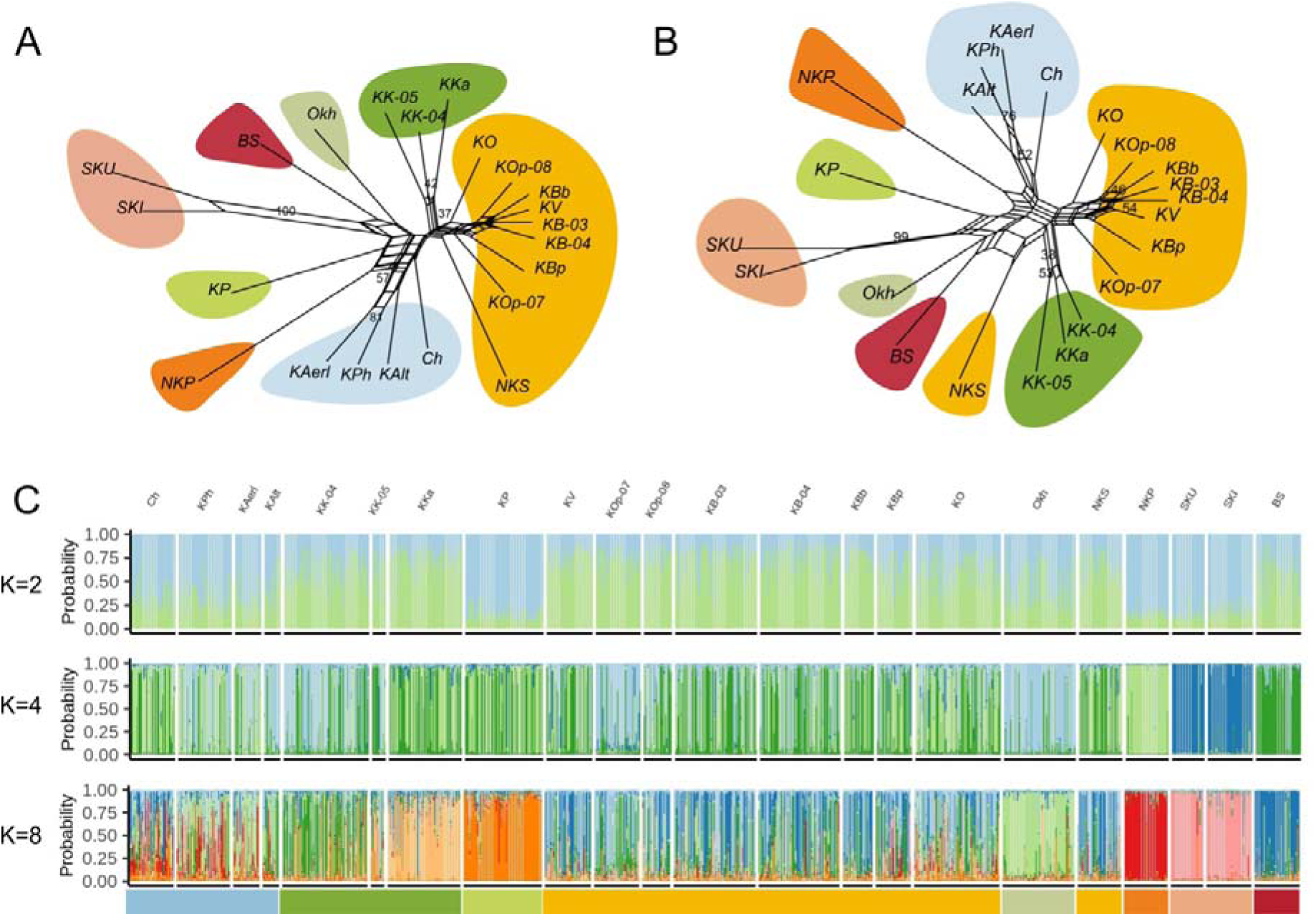
(**A**) Phylogenetic network build using the Neighbor-Net method and chord distances for samples of sockeye salmon from different watersheds of the Asian Pacific coast. Numbers in nodes are bootstrap indices (only values ≥25 are given); (**B**) The same for 27 putative neutral SNP loci; (С) An admixture analysis with STRUCTURE. The bar plots show the proportion of the genome assigned to each of the inferred clusters where each column represents an individual. Eight rounds of clustering were required to partition population structure down to region location. Color bars and areas are according to the color scheme in Fig. 1.

To enhance the accuracy of our phylogenetic reconstruction, we attempted to eliminate loci that may be subject to selection, potentially distorting the tree’s topology, and reconstruct the tree based on neutral substitutions. To achieve this, we applied a method for outlier loci detecting as proposed by Excoffier et al. for the hierarchical island model of populations (Excoffier et al., 2009). Specifically, *MHC2_190v2, MHC2_251v2, GPH-414, serpin, HGFA* (*p* < 0.01) and *GHII-2165, HpaI-99, U401-224, STC-410, ALDOB-135, GPDH* (*p* < 0.05) were identified as candidates for the directional selection, and *U504-141 & LEI-87* (*p* < 0.01) – as candidate loci for the balancing selection (Supplementary Figure S6). All the mentioned loci, as well as three mtSNPs, were subsequently excluded from the analysis. However, it’s noteworthy that the overall topology of the network remained largely unchanged, but the stability of some nodes in the already identified clades has increased, and the uncertainty in some basal nodes has increased significantly (Fig. 4B).

### Uncovering Population Structure by Bayesian Clustering of Samples

Based on the analysis conducted in STRUCTURE 2.3.4 and the determination of the optimal number of groups using STRUCTURE HARVESTER, the highest ΔK value was associated with eight clusters, the second highest peak corresponded to four clusters (Supplementary Figure S7). After the first round of clustering (K=2), island populations became isolated (except for the population of Shumshu Island), further clustering led to the consistent separation of the Okhota R. sample (K=3), Paramushir Island sample (K=4), Palana R. and Azabachye Lake samples (K=5) from a more or less homogeneous group of populations of Kamchatka and Chukotka (Fig. 4C). Further steps led to the separation of large clusters of South-West Kamchatka (including Shumshu Island) (SW) and North-Eastern cluster (NE). The distribution of samples across eight clusters corresponds to the subdivision of sockeye salmon in this region into nine distinct groups: SW, NE, and KR complexes, and individual populations of the Palana R., Okhota R., and three separate groupings of island populations of Bering Island, Paramushir Island and the South Kuril Islands (SK). The sockeye salmon population from Azabachye Lake was notably distinct from the other Kamchatka River samples, but did not exhibit differentiation when the MHC group loci were excluded from the analysis.

Spatial clustering analysis in GENELAND successfully identified eight distinct groups among Asian sockeye salmon populations, with a substantial overlap with the previously identified clusters in STRUCTURE (Fig. 5). The primary difference observed was the unification of two samples from watersheds within the Northern Kuril Islands into a single cluster.

**Fig. 5.**
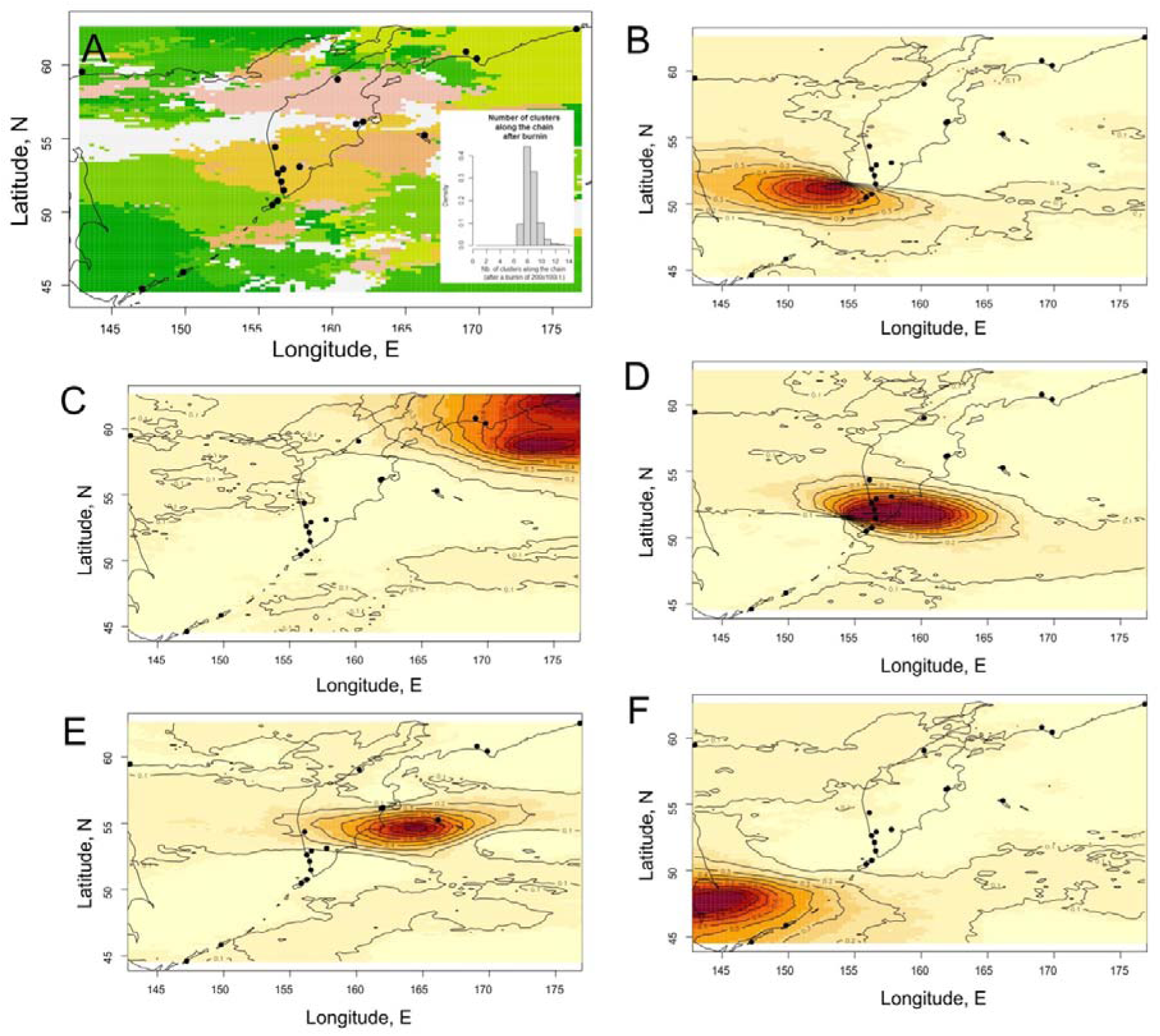
The results of the spatial Bayesian analysis implemented in GENELAND for Asian sockeye salmon: (**A**) A synthetic map of population membership in inferred clusters (spatial clusters marked by different colors) with posterior density distribution of the number of clusters (in the inset); (В**-F**) Panels showing space of the study area with the posterior probabilities belonging to some of inferred clusters for each pixel (dark red color reflects a higher posterior probability of membership to a cluster). Black dots represent the geographical position of sampled locations.

### Isolation by distance tests

To assess the load of gene flow in shaping territorial population complexes of sockeye salmon, we tested the hypothesis of isolation by distance separately for the Western and Eastern coasts of Kamchatka using more complete data from (Habicht et al., 2010).The Mantel test was employed to evaluate the correlation between matrices of genetic and geographic distances, both along Kamchatka coasts and within specific regions. The analysis revealed a significant correlation between geographic and genetic distances for both coasts of the peninsula, considering 27 neutral SNP loci (for East Kamchatka − *p* = 0.0005***, for West Kamchatka − *p* = 0.0341*). However, within the identified complexes, the tests were not significant (for populations of WK complex − *p* = 0.7679, for NE − *p* = 0.3471, for KR − *p* = 0.0834). These findings indicate a high level of population connectivity within complexes due to gene migration, with greater isolation observed between distant localities.

## Discussion

### Identifying Major Regional Complexes within Kamchatka peninsula

The analysis of sockeye salmon genetic differentiation within the studied range enables us to delineate regional population complexes associated with the most important reproduction areas of this species. The primary sockeye salmon populations in the Asian region reproduce within Kamchatka, where three distinct regional groups were identified: SW, NE, and KR complexes. Populations inhabiting watersheds along the North coast of the Sea of Okhotsk, specifically the Palana River and Okhota River, as well as island populations (excluding Shumshu Island population), were characterized by significant genetic differentiation from the so-called “core populations” of this species inhabiting Kamchatka rivers.

The establishment of sockeye salmon regional complexes is primarily associated with adaptations to their habitats in the same geographic area, the prevalence of life strategies adequate to local reproductive conditions, high connectivity through migrations, and is fundamentally determined by their common descent. Simultaneously, the main factors contributing to a specific range of adaptations within the complexes are related to the features of the spawning and freshwater growing watersheds, such as the topography, prevailing climatic conditions, its hydrobiological characteristics, aspects of anadromous migration and smoltification, as well as the coastal geomorphology that sets conditions for the early sea phase of juvenile life. The distinctions observed between the populations of West and East Kamchatka can be attributed to a complex set of adaptations specific to the climate and the landscape on the respective western and eastern coastlines of the peninsula, as well as prevailing types of spawning biotopes in the region. For instance, the length of the rivers on the west coast significantly exceeds that of the rivers of North-East Kamchatka. The lower and middle sections of the channels of most West Kamchatka rivers pass through low-lying swampy tundra and have a weak current (Pogodaev, 2013). In the majority of these watersheds, there are no deep lakes essential for the reproduction of this species, except for the Ozernaya and Palana rivers. Along the West Kamchatka, the coastline remains relatively flat, lacking significant bays or estuaries. Consequently, juvenile salmon from these rivers typically enter the open sea directly or briefly utilize estuary lakes or the estuarine zones of rivers for feeding. In contrast, the northeastern coast of Kamchatka indented by numerous bays and inlets. Juvenile salmon from this region first migrate to partially enclosed coastal waters, bays, and inlets before running to the open sea. This transitional phase may significantly influence their early marine survival (Pogodaev, 2013). Rivers on the northeast coast tend to be shorter and faster flowing. Most of them have lakes within their drainage basins (Supplementary Table S3). Within the populations inhabiting these regions, two distinct ecological-temporal forms are prevalent: an early form, primarily spawning in lakes, and a late form, predominantly reproducing in river bed. The distribution between these two forms within the populations is approximately equal (Shubkin & Bugaev, 2023). On the West Coast, the number of the early form is relatively low; the majority of the stocks are represented by the late, predominantly sea/river form. Given the higher migratory activity and lower homing of the sea/river ecitype, it is likely that moderate gene flow occurs between adjacent river basins, contributing to genetic uniformity within regional complexes (Beacham et al., 2006a). Consequently, the boundaries of such complexes can be determined by landscape zones and reproductive barriers. The gene migration restrictions can be caused by both the spatial distance or geographical barriers (such as seas, straits, mountain ridges, etc.), and ecological (specialization to the spawning ecotopes prevailing in a given area and subsequent isolation by adaptation) or temporary (separation by time of approaches to reproductive watersheds, anadromous migration and spawning sites) mechanisms of isolation.

The observed low level of population differentiation within the large Kamchatka complexes, SW and NE, can most likely be attributed to their common ancestral origin and the relatively short time that has passed since their divergence. It is commonly held that most populations of Asian sockeye salmon are relatively young, and their age does not exceed 10-12 thousand years (Chereshnev, 1998; Brykov et al., 2005), because its modern range largely coincides with the inferred area of the last Wisconsin glaciation in the North Pacific (maximum ∼26.5−19 thousand years ago, deglaciation ∼16,900−12,680 thousand years ago) (Braitseva & Evteeva, 1968; Velichko & Faustova, 1989). During the last glacial maximum, a substantial portion of the Western Kamchatka Lowland and the entire area of the Koryak Highlands were glaciated (Braitseva & Evteeva, 1968; Grosswald, 2009). This glaciation would have acted as a barrier preventing anadromous fish from accessing these watersheds. Previously, based on the results of an analysis of variability in the mtDNA control region, we determined that the entire Asian part of the sockeye salmon range represents a zone of secondary contact resulted from the rapid expansion of this species during the Holocene transgression, leading to the colonization of the majority of watersheds on the Kamchatka Peninsula by two distinct genetic lineages of sockeye salmon with differing origins (i.e., fish from different refugia that probably existed in the Beringia region (the territory of modern Alaska) and in the Kamchatka River basin) (Khrustaleva et al., 2020). These findings were confirmed by the results of admixture analysis in DAPC and in STRUCTURE, according to which all populations of these complexes had a “hybrid” origin, i.e. formed as a result of genetic admixture of at least two ancestral populations. It is evident that the contribution of North American populations to the gene pool of the northeastern complex is more substantial than that of the southwestern complex. This is further supported by the frequency distribution of combined mtSNP haplotypes in sockeye salmon, with a notable predominance of the GCC/CGG haplotype in the rivers of Northeastern Kamchatka and Chukotka (Khrustaleva, 2016). The same haplotype was dominant in American sockeye salmon populations from Norton Sound, Bristol Bay, Yukon and Kuskokwim rivers (Habicht et al., 2010).

### Metapopulation of the Kamchatka River Basin

Several authors in numerous studies have frequently proposed the existence of a major refugium within the Asian segment of its range, presumably located within the Kamchatka River paleobasin (Brykov et al., 2005; Beacham et al., 2006b; Khrustaleva et al., 2020). Presently, a large population of sockeye salmon is reproducing here, second only to the Kuril Lake population. The results of admixture analysis revealed the distinctiveness of the Azabachye Lake sample (formed a separate homogenous cluster), while sockeye salmon of the Kamchatka river basin had a hybrid origin (Fig. 3D, 4C) and contained an admixture of genotypes identical to those identified in Azabachye Lake. Genetic divergence of Azabachye and tributary sockeye salmon in the Kamchatka river basin has been repeatedly demonstrated in studies examining the polymorphism of microsatellite loci and mtDNA sequences (Brykov et al., 2005; Beacham et al., 2006b; Pilganchuk & Shpigalskaya, 2013; Pilganchuk et al., 2019; Khrustaleva et al., 2020). Additionally, Brykov (Brykov et al., 2005) postulated that this divergence can be attributed to the different origins of sockeye salmon in the upper and lower reaches of the river basin. Nevertheless, we posit that the differentiation of Azabachye sockeye salmon is likely associated with relatively recent demographic events. In the Azabachye Lake basin we examined the early sockeye salmon population, reproducing on the stream spawning grounds of a small river Bushuyka flowing into the lake. In the relatively small isolated population genetic drift usually occurs or it may have undergone fluctuations in its effective size in the recent past resulted in shifts in allelic frequencies. Positive bottleneck tests have been obtained for almost all samples from the Kamchatka River watershed including Azabachye sample (Table 1, Supplementary Table S3). In summary, we can deduce that the genetic characteristics unique to the metapopulation of Kamchatka river basin were caused by prolonged isolation of its part during the Upper Pleistocene glaciations within a large lake that existed in the middle reaches of the river (Bugaev & Kirichenko, 2008), subsequent rapid expansion into local and neighboring watersheds during the Holocene transgression, and, probably, secondary contact with adventive populations that colonized predominantly the lower part of the river basin after the glacier retreat (Brykov et al., 2005; Khrustaleva et al., 2020).

### Possible Factors Underlying the Divergence of North Sea of Okhotsk Populations

The high divergence observed in the Palana River population can result from the influence of directional selection. The selection is likely driven by the Palana River geographical location (subperiphery of the range) and the sockeye salmon biology peculiarities associated with its habitat conditions. The Palana River flows into the Sea of Okhotsk in the northern part of the peninsula at approximately 59°N. This region is characterized by a severe subarctic climate featuring long cold winters and short rainy summers. Palansky Lake, for example, remains ice-covered until late June, with ice cover fully restored by the end of October (Bugaev et al., 2002). The northern geographical location of the Palana River, far from the ocean growing areas, causes extensive migration of juvenile sockeye salmon along the coast of Western Kamchatka to reach the Pacific Ocean for feeding. This migratory pattern inevitably impacts their biological characteristics and population dynamics (Bugaev et al., 2002).

Furthermore, a distinctive feature of this population is that the spawning and freshwater feeding of juvenile sockeye salmon before migration to the sea are predominantly concentrated within the Palansky Lake. Here, in the vast majority of cases, sockeye salmon spend approximately 2 years (Bugaev et al., 2002), so all Palana sockeye salmon can be classified as a lake ecotype.

The Palana River estuary is hypertidal and characterized by significant spatiotemporal variability of all abiotic factors (primarily the level, speed and direction of water flow, salinity, temperature, turbidity) primarily driven by hyper-high tides, exceeding 10 meters. The tidal intrusion of salt water into the river’s channel can extend several kilometers upstream, leading to a strong reverse current. Juveniles migrating downstream have to expend more energy, in turn, high turbidity of water in the estuarine zone and active mixing of the water mass complicates the visual and olfactory orientation of migrants and the search for food. Consequently, the conditions for smoltification of sockeye salmon in this population can be classified as extreme. The survival of juveniles during the estuarine can only be supported by their high swimming activity, endurance, and readiness for the transition to saltwater (completeness of pre-catadromous transformation, achievement of a certain body size, corresponding physiological changes). In such a demanding environment, the mortality rate of underyearlings (0+) and yearlings (1+) is notably high.

Identifying the specific genes and mechanisms involved in these adaptive processes remains a complex challenge. The primary contribution to the third component that distinguishes this population from others is likely the genetic variations found in the *GPDH* (glycerol-3-phosphate dehydrogenase) and *RAG1* genes (Supplementary Fig. S3). The *GPDH2* SNP is localized in the CDS of the *GPDH* gene, encoding glycerol-3-phosphate dehydrogenase [NAD(+)], an enzyme involved in the Krebs cycle and cellular energy metabolism. A high level of its expression was detected in sockeye salmon smelt in British Columbia (Houde et al., 2019). The *RAG3-93* substitution is situated in the 3’ UTR of the recombination activating protein (*RAG1*) gene and may potentially be linked to adaptive loci within its coding region. The catalytic components of the *RAG* complex are involved in the activation of V-D-J recombination of immunoglobulin, which is a unique recombination mechanism that occurs only in developing lymphocytes at an early stage of B- and T-cell maturation and is responsible for the diversity of antibodies/immunoglobulins and T-cell receptors. The third SNP whose contribution to PC3 was quite significant was *VIM-569*. The *Vim* gene encodes vimentin, a protein found in intermediate filaments within connective tissues and other mesoderm-derived tissues. During vertebrate embryonic development, vimentin is expressed in cells with high migratory activity; postnatal expression of this gene is observed in fibroblasts, endothelial cells, lymphocytes and some specialized cells of the thymus and brain (Battaglia et al., 2018). Its main function is to maintain cellular and tissue integrity. Beyond this fundamental function, intermediate filaments play a crucial role in the intracellular distribution of organelles and proteins, subsequently influencing the functions of them (Minin & Moldaver, 2008). Several studies have demonstrated that the functioning of mitochondria within cells is influenced, in part, by the distribution of vimentin filaments (Herrmann & Aebi, 2000). Furthermore, two loci that stand out as potential candidates for directional selection in the Palana population are *ALDOB-135* and *GPH-414* (*p* < 0.005). These loci are located within the *ALDOB* and *GPH* genes, which encode Acyl-coenzyme A-binding protein and glycoprotein hormone alpha-subunit, respectively. The former is involved in lipid metabolism and the intracellular transport of acetyl-CoA, a central metabolite in the Krebbs cycle. The second is a common subunit of all pituitary hormones involved in the regulation of growth processes, puberty, the formation of breeding color and the resistance of fish to high temperatures.

Another perspective on the substantial divergence observed in the Palana River sample suggests that the reduced genetic diversity and a shift in gene frequencies could be attributed to a relatively recent reduction in effective population size, which was also confirmed by appropriate bottleneck tests. The sockeye salmon population in the Palana River experienced cyclical fluctuations similar to those observed in pink salmon (Bugaev, 2011). Bottleneck tests also support the hypothesis of effective population size decline in the nearest past. Thus, it becomes evident that the sockeye salmon population of Palana River has the special status among other sockeye salmon stocks of Western Kamchatka.

In the Okhota River, it appears that a somewhat different scenario is being realized. The neutrality tests did not reveal selection effects at any of the SNP loci in this sample. The notable reduction in genetic diversity within this sample suggests a recent decline in the effective population size rather than the influence of selection. The probability of a bottleneck in the Okhota River sockeye salmon population is confirmed by corresponding tests. Moreover, documented historical records support a decrease in the effective population size within this river basin. Thus, in the 1930s, more than 100 thousand sockeye salmon spawners passed through the Ueginsky lake system to spawn, and their total number in the river reached 1.5 million fish (Marchenko, 2022). In the 1960s − 1970s the number of sockeye salmon approaches has sharply decreased due to the significant impact of the Japanese drift-net fishery and the deterioration of conditions during the embryonic-larval period due to climatic cooling in the Sea of Okhotsk basin (Bugaev, 2011). For example, in the Ueginsky lakes in 1966, 1967 and 1968 about 20, 10 and 0.3 thousand spawners went to spawn, respectively, and in 1969–1971 – no more than 5–6 thousand fish (Nikulin, 1975). Furthermore, since 2016 total catch amounts increased, peaking at 479 tons in 2021 (Marchenko, 2022), but it is still subject to rather sharp fluctuations associated with the conditions of its natural reproduction.

### Island Populations: Distinctive Characteristics and Ecological Significance

The most substantial degree of genetic divergence, as determined through various tests and clustering methods, was observed between island and mainland populations. Likewise, significant distinctions were observed among island populations when compared to each other. The exception, apparently, is the sockeye salmon population of Shumshu Island, probably associated with the SW complex. Furthermore, within all the island populations examined, there was a pronounced reduction in genetic diversity, except for the sample from the Bettobu lake-river system (Shumshu Island). Notably, allelic diversity and expected heterozygosity were consistently lower in island populations than in mainland populations at all loci and neutral SNPs analyzed.

The pronounced genetic differences and decrease in genetic diversity of island populations can be explained by three primary categories of factors. The first category is associated with the geological history of this region and the patterns of the ichthyofauna of the Kuril and Commander ridges formation during periods of global climatic oscillations: long-term isolation in refugia and recurrent colonization of watersheds of the Kuril and Commander islands, founder effects or, which is also likely, extinction events due to greater amplitude of changes in environmental factors during global climate fluctuations on exposed land areas. Second, demographic factors such as small population sizes, effective number fluctuations, and a high degree of isolation will be considered. The distance of the islands from the center of the species’ range acts as a reproductive barrier, limiting gene flow not only between island and mainland populations but also between neighboring islands. Additionally, these populations are likely to have undergone a bottleneck event in the relatively recent past. Third, the observed genetic differences could reflect the development of local adaptations to specific environmental conditions within these island populations.

The similarity of samples from watersheds of the South Kuril Islands allows us to classify them as a single population complex. Evidently, this complex was formed under the influence of Pleistocene climatic cycles. The first Illinois glaciation covered all the islands of the Great Kuril Ridge. In the north, it was semi-glaciated, while the southern regions displayed characteristics of mountain-valley glaciation. The second glaciation, known as the Wisconsin, left minimal imprints on the central islands and primarily contributed to small moraines located at altitudes of 1000-1200 meters starting from Urup Island. Similar small moraines from the second glaciation were identified on Iturup Island, although they were absent further to the south. Consequently, the second glaciation extended over nearly all the islands, but in the north it was mountain-valley, and to the south it was close to cirque. During the Late Pleistocene regressions, it is presumed that Sakhalin, Hokkaido, Kunashir, the Lesser Kuril Ridge islands, and possibly Iturup were united as a single landmass, and had terrestrial connections with Primorye. (Gorshkov, 1967). It can be hypothesized that within the ice-free reservoirs of this extensive drained region of the continental shelf, populations were retained. These populations are likely descendants of the first wave of colonization who migrated from more northern territories and survived the most recent Wisconsin glaciation. The presence of Japanese kokanee populations isolated in the lakes of Hokkaido, located south of the primary species range, essentially serves as evidence of these glacial relic populations. During the Upper Pleistocene to early Holocene warming phase, temporarily exposed lands were flooded, populations dispersed and became isolated. Some of these populations subsequently formed relatively large stocks within suitable watersheds in the southern group of islands in the Kuril archipelago. It is probable that during the last Holocene transgression, these areas of the range were practically not subject to the expansion of colonists from more northern territories, as evidenced by mtDNA analysis data (Khrustaleva, 2016; Khrustaleva et al., 2020).

The islands in the northern region of the Kuril archipelago, specifically Paramushir and Shumshu, were physically connected to the Kamchatka Peninsula during the Late Pleistocene climatic cooling phases. However, while the origin of the Shumshu island population is confidently linked to the populations of the South-West Kamchatka complex, primarily due to its location along the migration routes of Ozernaya River sockeye salmon through the First and Second Kuril Strait (the distance between the mouth of the Bettobu River and the mouth of the Ozernaya River not exceeding 85 km), the population on Paramushir island likely has a more intricate history including the combined impact in its contemporary genetic diversity of the founder effect and genetic drift under long isolation. A similar scenario for the formation of genetic distinctiveness can be assumed for the Bering Island population. In this case, it’s likely that the primary factors influencing its genetic characteristics were the founder event and genetic drift. This assertion is supported by the analysis of mtSNP and the *CytB* gene sequences polymorphism, which revealed that the mass haplotype in Saranoye Lake was the same as in the Kamchatka River basin (Mineeva et al., 2015; Khrustaleva, 2016). This discovery suggests that the colonization of the Commander Islands occurred during the Holocene transgression from this nearby refugium. And it’s apparent that these islands were not heavily colonized by North American sockeye salmon for some reason. It is possible that the Kamchatka River basin was a fairly large center of expansion of this species on the Asian coast of the Pacific Ocean at the time. Alternatively, American alleles might have been subsequently eliminated due to genetic drift. In conclusion, the observed shifts in allelic frequencies and the decrease in genetic diversity in the Bering Island population may be the result of its descent from a limited number of individuals representing one or more ancestral gene pools. To comprehensively evaluate these hypotheses, more extensive studies are needed, including a broader sample selection from Asian, North American, and Japanese stocks, along with the analysis of different genetic markers, including sequences from various mtDNA segments.

The analysis conducted revealed a substantial loss of neutral genetic diversity in island forms of sockeye salmon compared to continental ones. Factors influencing within-population diversity can include bottlenecks, genetic drift, and inbreeding, all of which are dependent on the effective population size. Bottleneck tests across all island populations did not reveal a significant reduction in observed heterozygosity relative to expected, assuming equilibrium between mutations and drift. So, we can infer that effective population sizes have not experienced significant recent declines.

Microevolutionary processes within relatively small and isolated island populations are characterized by an increased influence of genetic drift, resulting in a reduction in genetic diversity. A marked decrease in allelic diversity and heterozygosity was observed across all loci as well as 27 neutral loci in all island populations, with the less pronounced effects seen in the sample from Shumshu Island and the greatest loss of diversity in Iturup Island population (Supplementary Figure 6). Notably, despite positive inbreeding coefficients (*F_IS_*) in most island populations, they were close to zero (as shown in Table 1). This suggests that mating conditions within these populations resemble panmix, with a relatively low occurrence of consanguineous crossing. Since directional selection also may lead to a reduction in genetic diversity, which may increase population viability and improve adaptation to the unique environment of a particular island, outlier SNPs have been identified when comparing island and mainland populations. Only two loci (*serpin* and *HGFA*) can be considered candidates for directional selection effects, which cannot explain the observed decrease in genetic diversity. Hence, it is likely that not inbreeding or natural selection, but genetic drift, which occurs under conditions of long-term isolation, played a predominant role in shaping the genetic characteristics of island populations and led to reduced genetic diversity and significant shifts in allelic frequencies, consequently influencing estimations of their genetic differentiation.

In conclusion, it is crucial to emphasize the significance of preserving biodiversity in unique island ecosystems. Many islands in the Kuril chain, including those we have examined, exhibit what is referred to as a “disharmony of the theriofauna,” characterized by a notable overabundance of predatory mammals in the species spectrum (Kostenko, 2002). On Iturup Island, for instance, the predator-to-prey ratio (comprising species such as fox, mink, sable, and bear as predators, and rat, red-gray vole, house mouse, and white hare as prey) stands at 1:1. In this context, predators are compelled to rely on supplementary trophic sources such as anadromous salmonids, which is an essential component of their diet on the islands. These findings underscore the necessity of preserving the intricate ecological balances that are essential for the overall health and stability of these unique ecosystems. Active commercial expansion and unregulated sockeye salmon fishery on the islands in conditions of limited ecological space can lead to the elimination of a number of species from island therio-complexes and increase instability of the species structure of the communities. The vulnerable balance of these island ecosystems is threatened by unsustainable exploitation and other negative anthropogenic impacts. Drawing from the findings we obtained in this and previous works, the southernmost populations of anadromous sockeye salmon in Asia (the populations inhabiting the lakes of the southern group islands of the Kuril ridge, along with the kokanee populations of Hokkaido) can be classified as glacial relicts. These populations possess unique genetic characteristics, exhibit distinct population dynamics, and are especially vulnerable to the impacts of global climate change and anthropogenic threats. Given their exceptional status, it is essential to pay special attention to them. Therefore, when developing a strategy for their sustainable commercial exploitation, it is important to establish systematic genetic and environmental monitoring programs. If there are indicators of a decline in their current condition, it is crucial to stop all fishing activities and initiate discussions regarding the implementation of protective measures of sockeye salmon within this region.

## Acknowledgments

Author gratefully acknowledges to Dr. J.E. Seeb (College of the Environment, University of Washington, Seattle, WA, USA) for his comprehensive assistance, provision of methods, and laboratory analysis management, as well as to all the employees of the Environmental Genomics Laboratory, Department of Hydrobiology and Fisheries, University of Washington, M.V. Shitova, Ph.D. (The Vavilov Institute of General Genetics of the Russian Academy of Sciences, VIGG RAS), and P.C. Afanasyev, Ph.D. (Russian Federal Research Institute of Fisheries and Oceanography, VNIRO), for their help in laboratory processing, N.V. Klovach, Dr.science (VNIRO), for sample collection management, and the employees of VNIRO, Kamchatka branch (KamchatNIRO), Pacific branch (TINRO), Sakhalin branch (SakhNIRO), and Magadan branch (MagadanNIRO) of the VNIRO, who took part in the samples collection.

## Supplementary Information

The online version contains supplementary material available at …

## Data Availability

The datasets generated and analyzed during the current study are available from the author on reasonable request.

## Ethical approval

This study was carried out in compliance with the Federal Law No. 498-FZ ‘On Responsible Handling of Animals and on Amending Certain Legislative Acts of the Russian Federation’ (17 December 2018). No ethical approval was required for fish provided dead by local fisheries.

