## Supplementary material for "Genetic Structuring and Conservation of Asian Sockeye Salmon: Identification of Regional Stock Complexes": Table S1

### Genetic Structuring and Conservation of Asian Sockeye Salmon: Identification of Regional Stock Complexes, Exploring Diversity, Origin, Adaptation, and Demography

**Hydrobiologia**


*Anastasia M. Khrustaleva*

Institute of Gene Biology Russian Academy of Sciences (FSBIS IGB RAS), Moscow, Russia


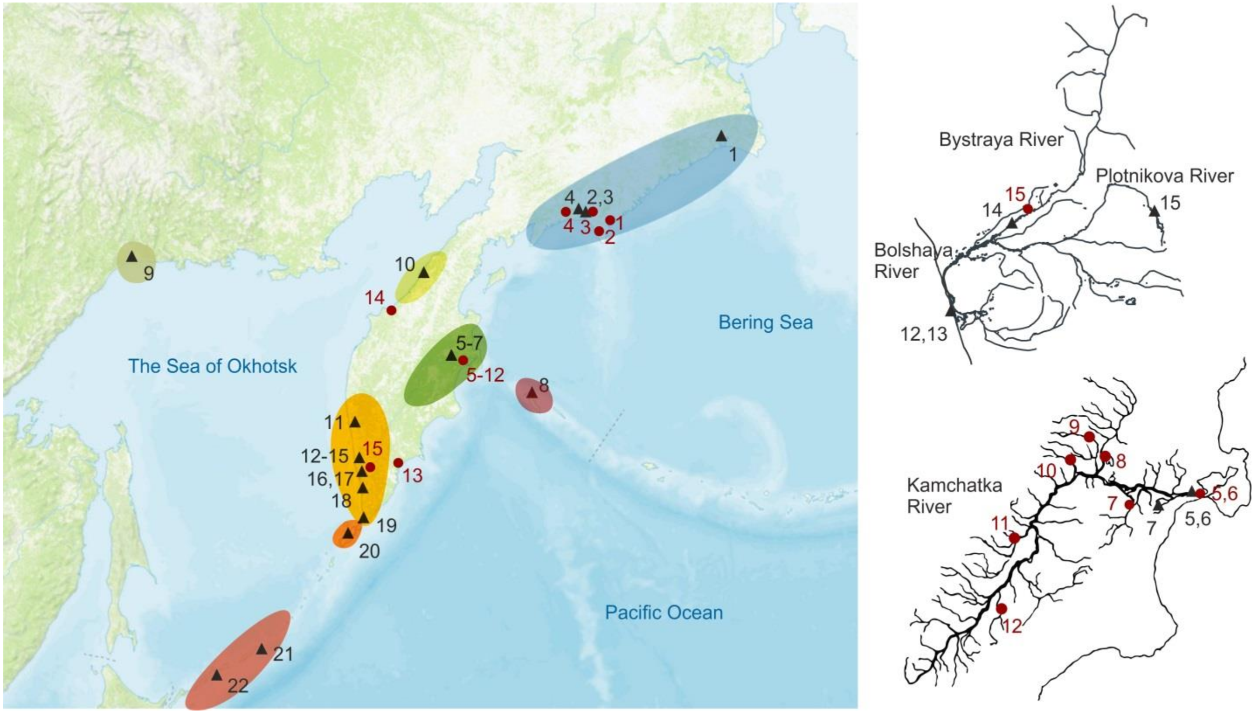


**Figure S1.** Schematic map of the study area with sampling points (triangles – our collection, red circles – data from Dr. Habicht et al. (Habicht et al., 2010)). The point's annotations are given in Table. 1 and Supplementary Table S1. Regional complexes of Asian sockeye salmon are marked with different colors.

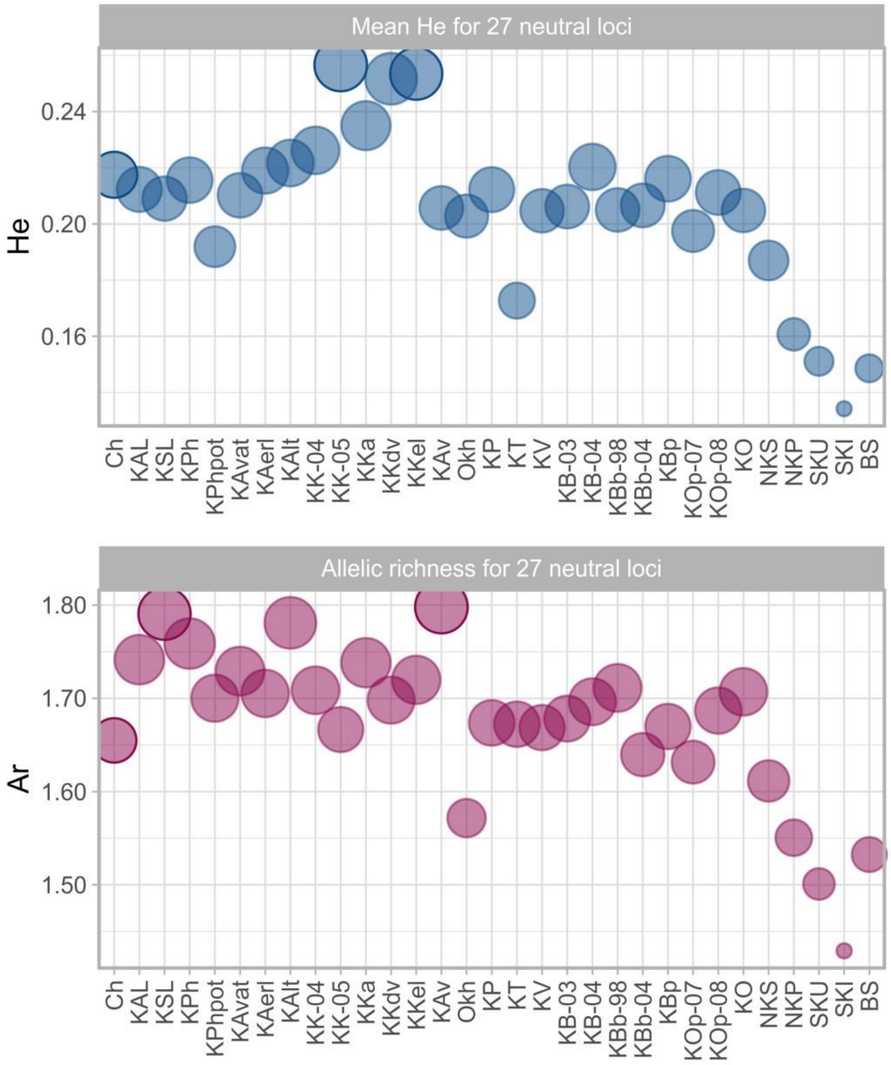


**Figure S2.** Mean expected heterozygosity and allelic richness for 27 neutral SNP loci in sockeye salmon populations along the Asian Cost of the Pacific Ocean.

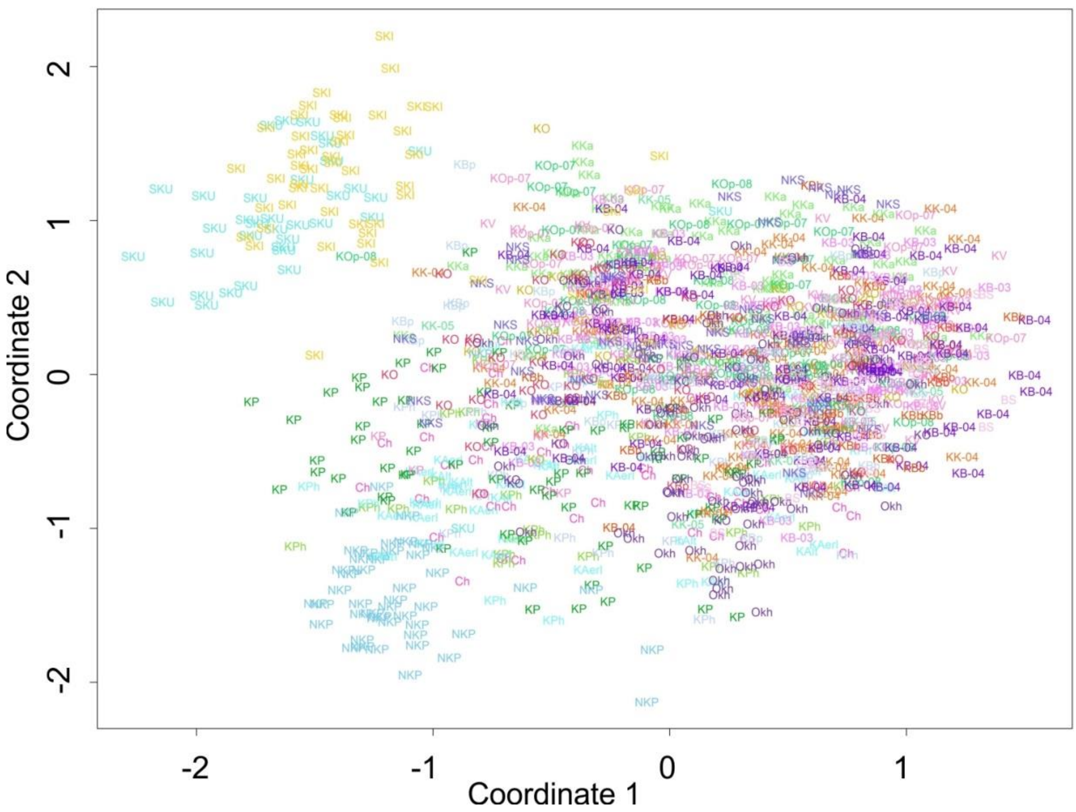


**Figure S3.** Individual multidimensional scaling plot (MDS or PCoA) based on Euclidian distances for sockeye salmon from the Asian coast of the Pacific Ocean. Samples are marked by their IDs and different colors.

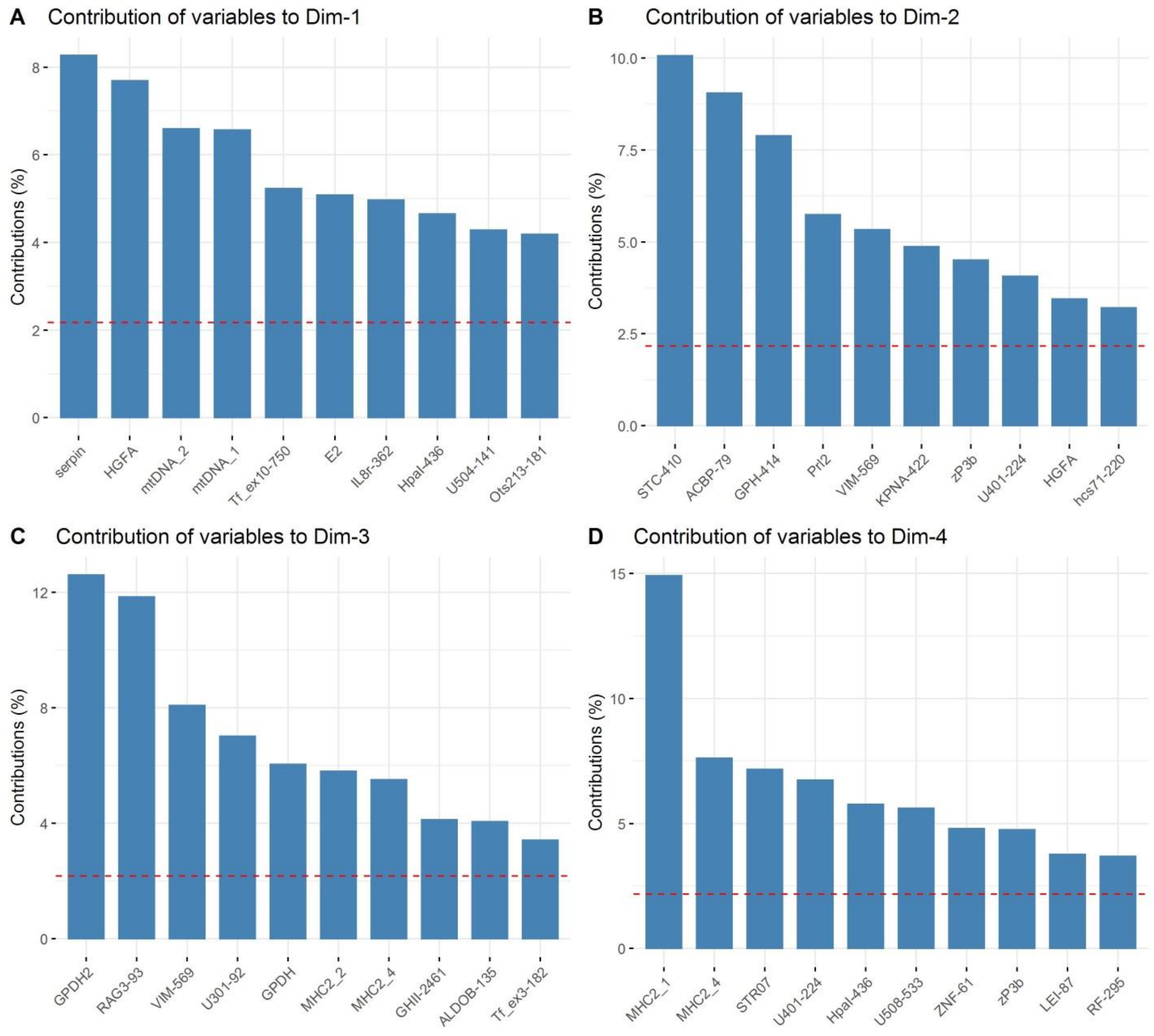


**Figure S4.** Principal component analysis (PCA) results: the contribution of loci to PC1-PC4. The top 10 contributors for each component are presented. The red dashed lines on the graphs indicate the expected average contribution; for a given component, a variable with a contribution larger than this cutoff could be considered important in contributing to the component.

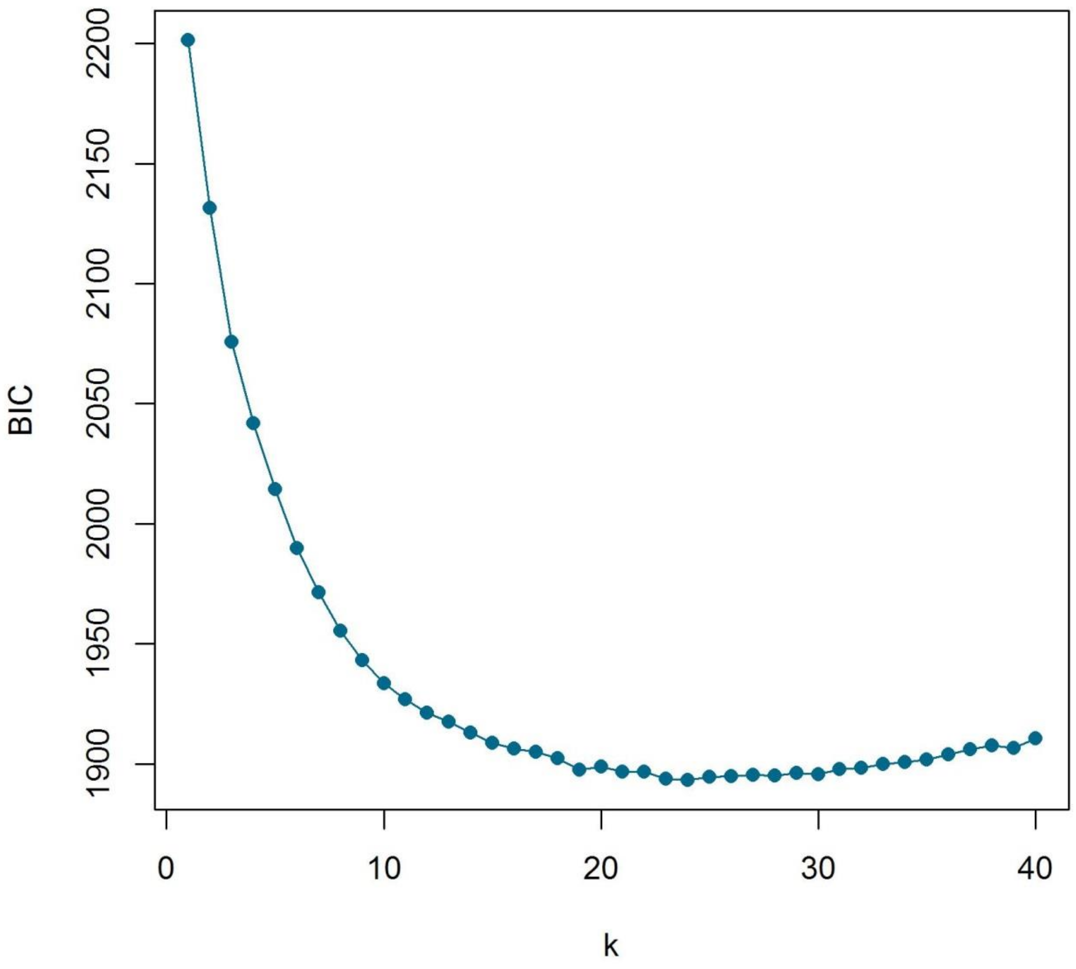


**Figure S5.** Bayesian information criterion (BIC) for number of clusters (k) in DAPC-analysis. The plot shows a rapid decrease in BIC until k = 8 clusters becomes the most likely value of K, and then a more gradual decline.

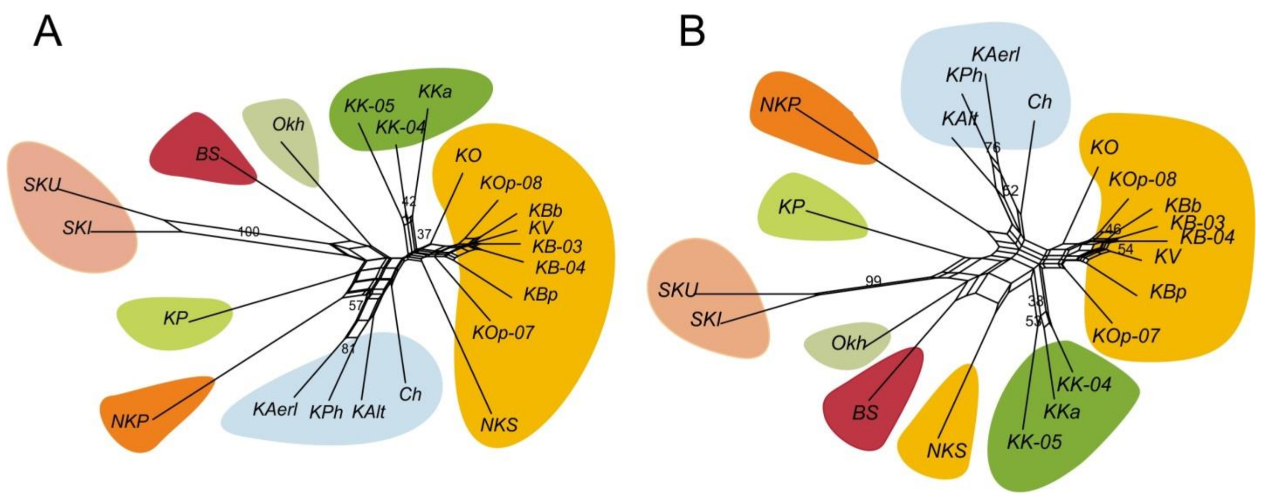


**Figure S6.** (A) Phylogenetic network build using the Neighbor-Net method and chord distances for samples of sockeye salmon from different watersheds of the Asian Pacific coast. Numbers in nodes are bootstrap indices (only values ≥25 are given); (B) The same for 27 putative neutral SNP loci

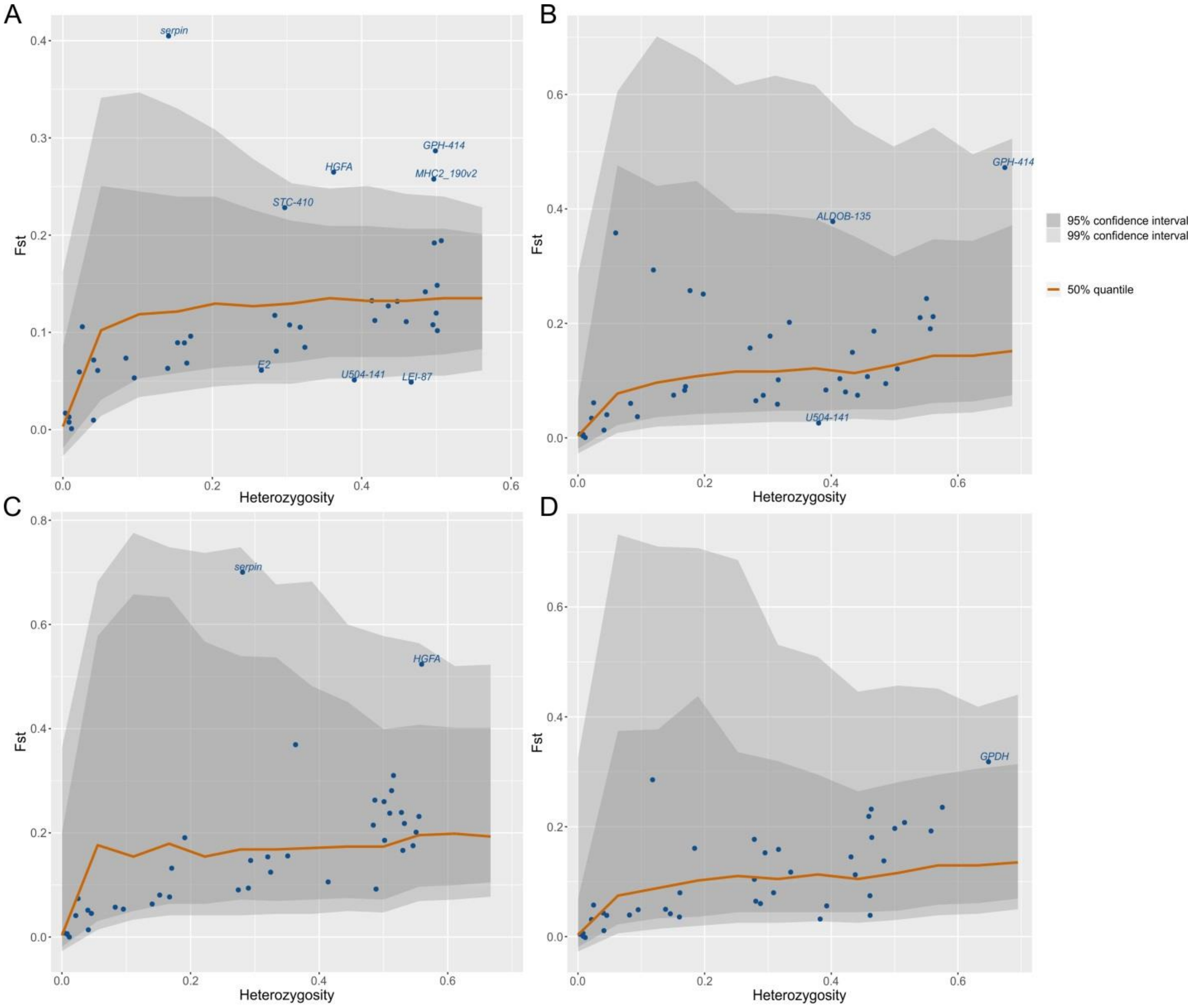


**Figure S7**. Examples of output images for outlier-SNP detection tests using Arlequin 3.5 for four combinations of samples: (A) – all samples, (B) – all samples vs KP sample, (C) – all continental samples vs all island populations, (D) – all samples vs BS sample. Loci falling above 5% (in the upper part of the graph) and below 1% (in the lower part of the graph) quantile limits were remooved as outliers. Here *serpin*, *HGFA*, *GPH-414*, *MHC2_190v2*, *STC-410*, *ALDOB-135*, *GPDH*, and *U504-141 & LEI-87* are considered as outliers.

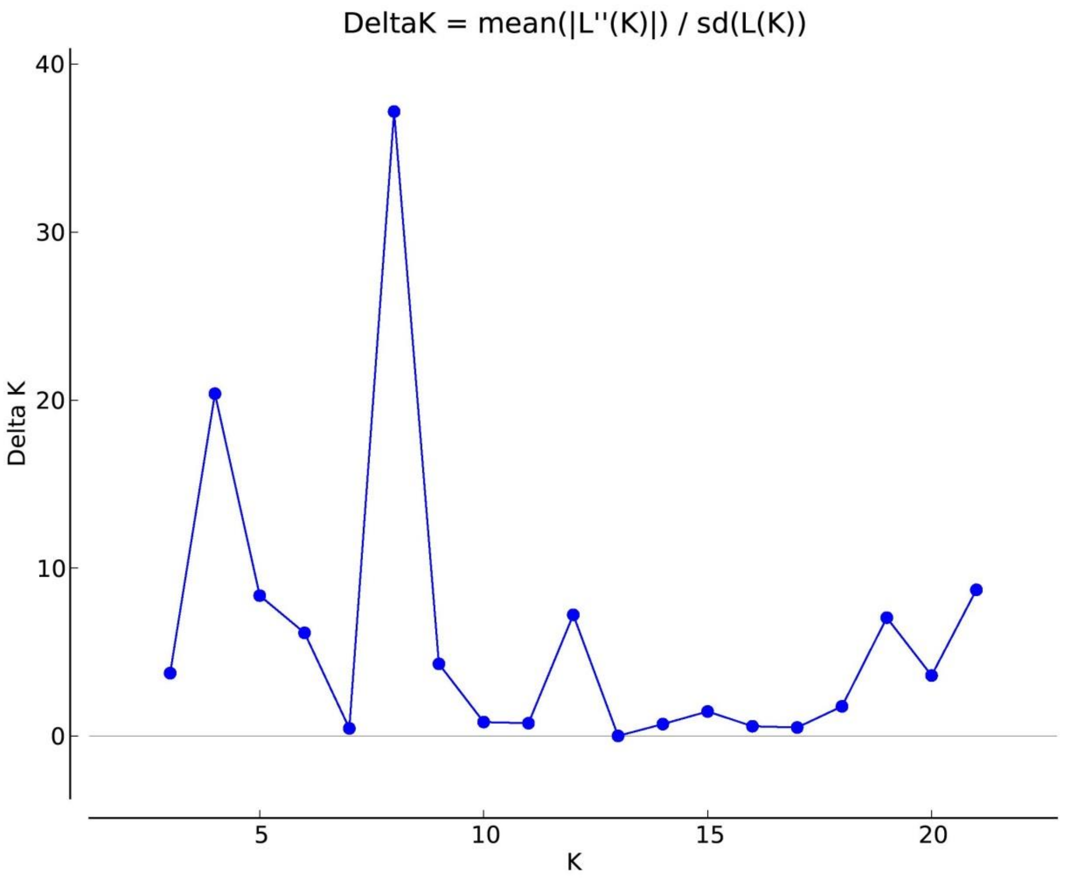


**Figure S8.** Delta K (ΔK) graph obtained by Structure Harvester with two maximums at K = 8 and 4.

**Table S1.** Sampling regions and locations, nursery lakes in the watershed, population IDs, and sampling methodology: date and place of catch, coordinates of a river mouth or a lake of catch, fishing gear and collector name if known.

| # | Region | Location | Nursery lakes | Pop. ID | Date of catch | Place of catch | Coordinates | Fishing gear | Collector |
|---|---|---|---|---|---|---|---|---|---|
| 1 | Chukotka, Navarin region | Meinypilgin lake-river system | Pikulneyskoye, High, Middle and Low Vaamochka lakes | Ch | 28.07.2004 | The creek between the Lower Vaamochka Lake and Vaamochka Lake | 62.540115°N, 176.813305°E | fixed nets, seine nets | E.V. Golub (Pacific branch of the VNIRO (“TINRO”)) |
| 2 | Kamchatka peninsula, Olyutor region | Apuka River, early run | Vatyt-Gythyn Lake | KAerl | 24.06.2008-25.06.2008 | The lower reaches | 60.418785°N, 169.849153°E | seine nets | V.I. Roy (VNIRO) |
| 3 | | Apuka River, late run | None | KAlt | 24.06.2008-25.06.2008 | The lower reaches | 60.418785°N, 169.849153°E | seine nets | V.I. Roy (VNIRO) |
| 4 | | Pakhacha River | Potat-Gythyn Lake | KPh | 17.06.2005-27.06.2005 | The lower reaches | 60.799973°N, 169.068806°E | seine nets | E.D. Pavlov (VNIRO) |
| 5 | Kamchatka peninsula, East coast | Kamchatka River, late run | Azabachje Lake, Dvu-Yurtochnoye Lake, Kursin Lake, Kamakov lowland lakes | KK-04 | 29.06.2004-09.07.2004 | Downstream, 5 and 30 km from the mouth | 56.192783°N, 161.997488°E | seine nets | A.M. Khrustaleva (IGB RAS) |
| 6 | | Kamchatka River, early run | | KK-05 | 14.06.2005 | Downstream, 5 and 30 km from the mouth | 56.192783°N, 161.997488°E | seine nets | V.I. Roy (VNIRO) |
| 7 | | Azabachje Lake, Bushuyka River | Azabachje Lake | KKa | 03.07.2004, 13.07.2004 | Bushuyka River outfall (tributary of Azabachje Lake) | 56.119205°N, 161.854666°E | seine nets | A.M. Khrustaleva (IGB RAS), V.F. Bugaev (Kamchatka branch of the VNIRO (“KamchatNIRO”)) |
| 8 | Commander Islands | Bering Island, Sarannoye Lake | Sarannoye Lake | BS | 7.2008 | Sarannoye Lake | 55.271646°N, 166.146979°E | seine nets | no data |
| 9 | Continental coast of the Sea of Okhotsk | Okhota River | Ueginsky Lakes | Okh | 22.07.2004 | The lower reaches | 59.469398°N, 142.952526°E | seine nets | Magadan branch of the VNIRO (“MagadanNIRO”) |
| 10 | Kamchatka peninsula, North-West | Palana River | Plansky Lake | KP | 10.07.2003-21.07.2003 | Downstream, 10 km from the mouth | 59.035209°N, 160.213639°E | gill nets, seine nets | A.M. Khrustaleva (IGB RAS) |
| 11 | Kamchatka peninsula, South-West | Bolshaya Vorovskaya River | Vorovsky Lake, No name Lake, Kapovoye Lake | KV | 17.07.2007-27.07.2007 | The lower reaches | 54.359467°N, 156.095038°E | seine nets | Kamchatka branch of the VNIRO (“KamchatNIRO”) |
| 12 | | Bolshaya River | Nachikinsky Lake, Sokotch Lake | KB-03 | 23.07.2003-30.07.2003 | Downstream, 5-10 km from the mouth | 52.628162°N, 156.26196°E | seine nets | A.M. Khrustaleva (IGB RAS) |

| | | | | | | | | | |
|---|---|---|---|---|---|---|---|---|---|
| 13 | | Bolshaya River | | KB-04 | 11.08.2004-20.08.2004 | The lower reaches | 52.628162°N, 156.26196°E | seine nets | R.A. Zboev, I.N. Kireev (Kamchatka branch of the VNIRO (“KamchatNIRO”)) |
| 14 | | Bolshaya River drainage, Bistraya River | None | KBb | 20.07.2004-12.08.2004 | The lower course of Bystraya River, 10 km from the Karymay settlement | 52.92024°N, 156.609568°E | minnow seine | E.V. Yesin (VNIRO) |
| 15 | | Bolshaya River drainage, Plotnikova River | Nachikinsky Lake, Sokotch Lake | KBp | 09.08.2004-12.082004 | Upper reach of Plotnikov River, 10 km from Nachikinskoe Lake | 53.10024°N, 157.756895°E | minnow seine | E.V. Yesin (VNIRO) |
| 16 | | Opala River | Opalinsky Lake | KOp-07 | 01.07.2007 | The lower reaches | 52.131653°N, 156.476848°E | seine nets | A.I. Manukhov (VNIRO) |
| 17 | | Opala River | Opalinsky Lake | KOp-08 | 17.07.2008-26.08.2009 | The lower reaches | 52.131653°N, 156.476848°E | seine nets | S.A. Belorusceva (VNIRO) |
| 18 | | Ozernaya River | Kurilsky Lake, Etamynk Lake | KO | 04.08.2003-07.08.2003 | The lower reaches | 51.484258°N, 156.568087°E | seine nets | A.M. Khrustaleva (IGB RAS) |
| 19 | North Kuril Islands | Shumshu Island, Bettobu Lake | Bettobu Lake | NKS | 05.08.2008 | Ostrognaya Ryver (tributary of Bettobu Lake) | 50.751485°N, 156.264268°E | seine nets | K.V. Chudanov (VNIRO) |
| 20 | | Paramushir Island, Glukhoye Lake | Glukhoye Lake | NKP | 07.07.2008-13.07.2008 | Shumnaya Ryver (flows into Glukhoye Lake) | 50.488928°N, 155.847357°E | seine nets | no data |
| 21 | South Kuril Islands | Urup Island, Tokotan Lake | Tokotan Lake | SKU | 07.2008-08.2008 | Tokotan Lake | 45.858739°N, 149.799329°E | seine nets | Sakhalin branch of the VNIRO (“SakhNIRO”) |
| 22 | | Iturup Island, Krasivoye Lake | Krasivoye Lake | SKI | 01.10.2006 | Krasivoye Lake | 44.624539°N, 147.208965°E | seine nets | Sakhalin branch of the VNIRO (“SakhNIRO”) |

**Table S2:** Characteristics of 45 SNP loci. *He* – mean expected heterosigocity, *Ho* – mean observed heterosigocity, $n_a$ – mean allele count per locus, $F_{ST}$ − fixation index by locus.

| # | Locus name | GenBank ID | Putative location | SNP position | Substi-tution | Description | $H_e(SD)$ | $H_o(SD)$ | $n_a(SD)$ | $F_{ST}$ |
|---|---|---|---|---|---|---|---|---|---|---|
| 1 | *One_ACBP-79* | DQ386287 | acyl-coenzyme A-binding protein (ACBP) gene | 79 | A/G | mRNA | 0.39(0.14) | 0.36(0.13) | 1.91(0.29) | 0.132 |
| 2 | *One_ALDOB-135* | DQ386280 | aldolase B (ALDOB) gene, partial sequence | 135 | G/A | noncoding region | 0.22(0.15) | 0.21(0.13) | 1.86(0.35) | 0.117 |
| 3 | *One_CO1* | AY353070 | haplotype TAGG DNA cytochrome oxidase I-like gene, partial sequence; mitochondrial | 7061 | T/C | synonymous | haploid | | 1.96(0.2) | – |
| 4 | *One_ctgf-301* | DQ386288 | connective tissue growth factor (CTGF) gene | 287 | G/T | mRNA | 0.01(0.02) | 0.01(0.02) | 1.23(0.43) | 0.013 |
| 5 | *One_Cytb_17* | AY353063 | GT DNA cytochrome b-like gene, partial sequence; mitochondrial | 16162 | G/A | synonymous | haploid | | 1.17(0.38) | – |
| 6 | *One_Cytb_26* | AY353064 | GC DNA cytochrome b-like gene, partial sequence; mitochondrial | 16168 | T/C | synonymous | haploid | | 1.96(0.2) | – |
| 7 | *One_E2* | DQ025695 | One.E2.31.36 genomic sequence similar to type II keratin E2 | 65 | A/G | mRNA | 0.25(0.11) | 0.26(0.14) | 2(0) | 0.061 |
| 8 | *One_GHII-2461* | U14535.1 | type-2 growth hormone gene, complete cds | 43 | T/A | intron | 0.14(0.14) | 0.15(0.15) | 1.82(0.39) | 0.089 |
| 9 | *One_GPDH* | DQ025723 | One.GPDH.40.61 genomic sequence similar to glycerol-3-phosphate dehydrogenase (GPDH) | 187 | C/G | mRNA | 0.44(0.11) | 0.44(0.14) | 2(0) | 0.108 |
| 10 | *One_GPDH2* | DQ025724 | One.GPDH.56.48 genomic sequence similar to glycerol-3-phosphate dehydrogenase (GPDH) | 201 | C/T | mRNA | 0.13(0.11) | 0.12(0.12) | 1.95(0.21) | 0.063 |
| 11 | *One_GPH-414* | DQ386289 | glycoprotein hormone alpha-subunit (GPH) gene | 414 | T/C | intron | 0.36(0.13) | 0.35(0.14) | 2(0) | 0.287 |
| 12 | *One_hcs71-220* | DQ386293 | major heat shock protein-like protein (HSC71) gene, partial sequence | 220 | A/C | intron | 0.4(0.13) | 0.39(0.13) | 2(0) | 0.111 |
| 13 | *One_HGFA* | DQ025719 | One.HGFA.22.46 genomic sequence similar to hepatocyte growth factor activator/GRAAL (HGFA) | 49 | A/T | unknown | 0.26(0.14) | 0.24(0.14) | 1.95(0.21) | 0.265 |
| 14 | *One_HpaI-436* | DQ386294 | One-436 HpaI repeat element-like sequence similar to HpaI repeat element | 79 | A/T | unknown | 0.42(0.08) | 0.39(0.09) | 2(0) | 0.148 |
| 15 | *One_HpaI-99* | DQ386281 | clone One-99 HpaI repeat element-like sequence | 99 | C/T | unknown | 0.08(0.1) | 0.08(0.1) | 1.5(0.51) | 0.073 |
| 16 | *One_IL8r-362* | FM206384, GU570948.1 | Oncorhynchus mykiss partial il-8 gene for interleukin 8, promoter region (interleukin-8 receptors), isolate 98JN-020 chemokine receptor (IL8R) gene | 362 | C/T | promoter region | 0.29(0.1) | 0.27(0.13) | 2(0) | 0.108 |
| 17 | *One_KPNA-422* | DQ386282 | karyopherin alpha 2 (KPNA2) gene | 422 | A/G | unknown | 0.34(0.14) | 0.34(0.14) | 1.95(0.21) | 0.133 |
| 18 | *One_LEI-87* | DQ386279 | leukocyte elastase inhibitor (LEI) gene | 87 | A/G | unknown | 0.45(0.11) | 0.47(0.13) | 2(0) | 0.049 |
| 19 | *One_MARCKS-241* | GU570949.1 | myristoylated alanine-rich protein kinase (MARCKS) gene | 241 | A/T | unknown | 0.01(0.01) | 0.01(0.01) | 1.36(0.49) | 0.001 |
| 20 | *One_MHC2_190v2* | AY386256 | major histocompatibility complex class II B1 (Onne-DAB) gene, Onne-DAB-3 allele, partial cds | 190 | T/G | exon | 0.37(0.13) | 0.33(0.16) | 1.97(0.12) | 0.258 |

| | | | | | | | | | | |
|---|---|---|---|---|---|---|---|---|---|---|
| 21 | *One_MHC2_251v2* | AY386257 | major histocompatibility complex class II B1 (Onne-DAB) gene, Onne-DAB-4 allele, partial cds | 251 | C/T | intron | 0.41(0.12) | 0.35(0.17) | 1.98(0.1) | 0.192 |
| 22 | *One_Ots213-181* | DQ386285 | clone Ots213 genomic sequence | 220 | T/G | unknown | 0.16(0.15) | 0.15(0.14) | 1.86(0.35) | 0.096 |
| 23 | *One_p53-576* | DQ386284 | p53 tumour suppression gene, partial sequence | 534 | A/C | unknown | monomorphic | | 1(0) | – |
| 24 | *One_pIns-107* | DQ025686 | One.ins.10.30 genomic sequence similar to insulin gene | 107 | C/T | unknown | 0.46(0.06) | 0.43(0.08) | 2(0) | 0.102 |
| 25 | *One_Prl2* | AY353071 | haplotype G prolactin II gene, partial cds (coding sequence) | 187 | G/T | coding | 0.44(0.1) | 0.44(0.13) | 2(0) | 0.12 |
| 26 | *One_RAG1-103* | DQ386290 | recombination activating protein (RAG1) gene | 103 | A/T | unknown | monomorphic | | 1(0) | – |
| 27 | *One_RAG3-93* | DQ386291 | recombination activating protein (RAG1) gene | 93 | C/T | mRNA | 0.03(0.06) | 0.03(0.05) | 1.5(0.51) | 0.072 |
| 28 | *One_RF-112* | AB435387 | 12-RFa mRNA for 12-RF amide peptide, complete cds | 112 | A/G | exon, synonymous | 0.26(0.13) | 0.26(0.13) | 2(0) | 0.081 |
| 29 | *One_RF-295* | AB435387 | 12-RFa mRNA for 12-RF amide peptide, complete cds | 259 | A/T | exon | 0.04(0.08) | 0.04(0.08) | 1.5(0.51) | 0.061 |
| 30 | *One_RH2op-395* | DQ386277 | RH2 opsin (RH2op) gene, partial sequence | 395 | T/G | unknown | 0.04(0.04) | 0.04(0.04) | 1.73(0.46) | 0.01 |
| 31 | *One_serpin* | DQ025707 | clone One.serpin.50.38 genomic sequence similar to SERine proteinase Inhibitors (serpin) | 75 | T/G | unknown | 0.09(0.13) | 0.09(0.14) | 1.73(0.46) | 0.405 |
| 32 | *One_STC-410* | DQ386278 | stanniocalicin (STC) gene | 410 | C/T | intron | 0.21(0.16) | 0.2(0.16) | 1.95(0.21) | 0.228 |
| 33 | *One_STR07* | DQ386286 | clone STR07 genomic sequence | 182 | C/G | unknown | 0.29(0.17) | 0.27(0.16) | 1.91(0.29) | 0.085 |
| 34 | *One_Tf_ex10-750* | AH015399.2 | transferrin (Tf) gene | 750 | A/G | exon | 0.42(0.13) | 0.39(0.15) | 2(0) | 0.142 |
| 35 | *One_Tf_ex3-182* | AH015399.2 | transferrin (Tf) gene | 182 | A/G | intron | 0.01(0.02) | 0(0.01) | 1.23(0.43) | 0.008 |
| 36 | *One_U301-92* | DQ267490 | FK506-binding protein 12-like (FKBP12) gene | 92 | G/T | unknown | 0.15(0.13) | 0.14(0.14) | 1.91(0.29) | 0.068 |
| 37 | *One_U401-224* | GU570950.1 | unknown | 224 | A/C | unknown | 0.36(0.1) | 0.34(0.1) | 2(0) | 0.112 |
| 38 | *One_U404-229* | GU570951.1 | unknown | 229 | C/T | unknown | 0.03(0.07) | 0.03(0.07) | 1.27(0.46) | 0.106 |
| 39 | *One_U502-167* | GU570952.1 | unknown | 167 | A/G | unknown | 0(0.02) | 0(0.02) | 1.09(0.29) | 0.017 |
| 40 | *One_U503-170* | GU570953.1 | unknown | 170 | G/T | unknown | 0.28(0.16) | 0.28(0.17) | 1.91(0.29) | 0.105 |
| 41 | *One_U504-141* | GU570954.1 | unknown | 141 | A/C | unknown | 0.36(0.14) | 0.36(0.14) | 1.95(0.21) | 0.051 |
| 42 | *One_U508-533* | GU570955.1 | unknown | 162 | C/T | unknown | 0.1(0.1) | 0.08(0.09) | 1.77(0.43) | 0.053 |
| 43 | *One_VIM-569* | DQ386292 | vimentin (VIM) gene | 563 | A/G | unknown | 0.14(0.13) | 0.15(0.14) | 1.91(0.29) | 0.089 |
| 44 | *One_ZNF-61* | BT057144 | ER lumen protein-retaining receptor 2 | 61 | C/A | mRNA | 0.37(0.13) | 0.34(0.13) | 2(0) | 0.127 |
| 45 | *One_zP3b* | DQ025739 | Oncorhynchus mykiss C-C chemokine receptor type 9-like | 49 | A/C | mRNA | 0.03(0.06) | 0.03(0.07) | 1.27(0.46) | 0.059 |

**Table S3.** Samples characteristics, regions and locations, population IDs, date of catch, coordinates, and summary statistics for 40 SNP loci according to the data from (Habicht et al., 2010): mean expected (*He*) heterozygosities, allelic richness (*Ar*), and the results of sign test ($p_{sign}$), standardized differences test ($p_{stdv}$) and two-tail Wilcoxon sign-rank tests ($p_W$) for heterozygote excess, * − $p < 0.05$, ** − $p < 0.01$, *** − $p < 0.001$.

| # | Region | Location | Pop. ID | Date of catch | Coordinates | *n* | *Ar(SD)* | *He(SD)* | $p_{sign}$ | $p_{stdv}$ | $p_W$ |
|---|---|---|---|---|---|---|---|---|---|---|---|
| 1 | Kamchatka peninsula, Olyutor region | Severnaya Lagoon | KSL | 6/26/2002 | 60.5°N, 170.62°E | 98 | 1.93(0.33) | 0.21(0.18) | 0.107 | 0.035* | 0.134 |
| 2 | | Anana Lagoon | KAL | 6/24/2002 | 60.04°N, 170.22°E | 80 | 1.78(0.42) | 0.21(0.19) | 0.008** | 0.002** | 0.038* |
| 3 | | Apuka River, Vatit Lake | KAvat | 8/7/2002 | 60.67°N, 170.25°E | 51 | 1.78(0.42) | 0.21(0.18) | 0.004** | 0.012* | 0.05 |
| 4 | | Pakhacha River, Potat Lake | KPhpot | 7/29/2001 | 60.67°N, 167.6°E | 50 | 1.74(0.43) | 0.19(0.18) | 0.007** | 0.026* | 0.064 |
| 5 | Kamchatka River basin | Kamchatka River, late run | KKlt-98 | 7/21/1998 | 56.23°N, 162.5°E | 100 | 1.78(0.42) | 0.21(0.19) | 0.008** | 0.002** | 0.024* |
| 6 | | Kamchatka River, early run | KKerl-98 | 6/1/1998 | 56.23°N, 162.5°E | 78 | 1.74(0.44) | 0.19(0.17) | 0.048* | 0.01* | 0.04* |
| 7 | | Hapiza River | KKhap | 9/2/1998 | 56.2°N, 161.25°E | 146 | 1.74(0.44) | 0.23(0.19) | 0.001** | 0*** | 0.002** |
| 8 | | Elovka River | KKel | 1995 | 56.58°N, 160.75°E | 109 | 1.78(0.42) | 0.25(0.2) | 0*** | 0*** | 0*** |
| 9 | | Dvu 'Yurta River | KKdv | 1995 | 56.55°N, 160.13°E | 88 | 1.78(0.43) | 0.25(0.21) | 0*** | 0*** | 0.001** |
| 10 | | Belaya River | KKbel | 1995 | 56.43°N, 160.35°E | 81 | 1.85(0.37) | 0.23(0.19) | 0.001** | 0.002** | 0.02* |
| 11 | | Kozireuka River | KKkoz | 1994 | 56.03°N, 159.78°E | 40 | 1.74(0.44) | 0.22(0.18) | 0.008** | 0.003** | 0.006** |
| 12 | | Kitilgina River, early run | KKkit | 6/29/1998 | 55.07°N, 159.13°E | 28 | 1.78(0.42) | 0.2(0.18) | 0.234 | 0.086 | 0.338 |
| 13 | Kamchatka peninsula, South-East | Avacha Bay | KAv | 2002 | 53.03°N, 158.44°E | 60 | 1.85(0.35) | 0.21(0.18) | 0.071 | 0.041* | 0.134 |
| 14 | Kamchatka peninsula, North-West | Tigil River | KT | 6/18/2002 | 58°N, 158.27°E | 107 | 1.81(0.41) | 0.17(0.18) | 0.035* | 0.089 | 0.388 |
| 15 | Kamchatka peninsula, South-West | Bistraya River | KBb-98 | 8/16/1998 | 53.35°N, 157.47°E | 56 | 1.74(0.44) | 0.2(0.19) | 0.055 | 0.005** | 0.036* |

**Table S4.** Deviations from Hardy–Weinberg equilibrium in samples revealed at some loci after FDR-correction: *p*-value – the exact HWE test results, W&C and R&H − two estimates of $F_{IS}$, Weir & Cockerham's (1984) estimate and Robertson & Hill's (1984) estimate.

| Sample | Locus | *p*-value | W&C | R&H |
|---|---|---|---|---|
| KK-04 | *MHC2_190v2* | 0.0003 | 0.6515 | 0.6578 |
| KK-04 | *MHC2_251v2* | 0 | 0.9547 | 0.9646 |
| KB-03 | *ZNF-61* | 0.0004 | 0.3857 | 0.3886 |
| KB-04 | *ZNF-61* | 0.0002 | 0.3989 | 0.4021 |
| KB-04 | *Tf_ex3-182* | 0.0001 | 0.4929 | 0.497 |
| KBp | *MHC2_251v2* | 0.0003 | 0.5979 | 0.6104 |
| Okh | *GPDH2* | 0.0001 | 0.492 | 0.4966 |
| Okh | *GPH-414* | 0.0001 | 0.4293 | 0.4332 |
